# Activated Interferon Signaling Suppresses Age-Dependent Liver Cancer

**DOI:** 10.1101/2024.07.31.606057

**Authors:** Aaron P. Havas, Kathryn Lande, Adarsh Rajesh, K. Garrett Evensen, Siva Karthik Varanasi, Linshan Shang, Elizabeth Schmidt, Jin Lee, Kenneth Kim, Marcos Garcia Teneche, Filipe Hoffmann, Michael LaPorte, Andrew Davis, Abby Grier, Julie A. Reisz, Kevin Tharp, Armin Gandhi, Xue Lei, Jessica Proulx, Karl N. Miller, Alessandra Sacco, Gerald S. Shadel, Laura Niedernhofer, Gen-Sheng Feng, Angelo D’Alessandro, Susan Kaech, April Williams, Peter D. Adams

## Abstract

Age is a major risk factor for liver cancer, as is the case for most adult human cancers. However, the underlying mechanisms are not well defined. A better understanding of the role of aging in liver and other cancers can facilitate approaches for risk assessment, early detection and prevention. We hypothesize that age-driven changes render aged liver more sensitive to oncogenic stress and hence tumorigenesis. To investigate how the liver changes with age, we documented the immune profile, transcriptome and epigenome of healthy livers from both young and aged mice, revealing pronounced alterations with aging. Notably, in aged hepatocytes, we identified heightened interferon (IFN) signaling, as well as simultaneous tumor suppressor and oncogene signaling at both bulk and single cell level, suggestive of an aged liver that is poised for neoplasia. To challenge this seemingly poised state, we employed adeno-associated virus (AAV)-mediated expression of a c-Myc oncogene in young and aged mouse liver hepatocytes *in vivo*. Analysis of aged hepatocytes expressing c-Myc revealed further elevated expression of IFN Stimulated Genes (ISGs). This ISG upregulation was evident in multiple models of oncogenic stress and transformation in older mice and also observed in aged humans with Metabolic dysfunction-Associated Steatohepatitis (MASH). We determined that Stat1 is both necessary and sufficient for the age specific elevated ISG expression in old wild type mice. Remarkably, inhibiting Jak/Stat signaling alongside ectopic c-Myc expression led to high-grade hepatocyte dysplasia and tumor formation, selectively in aged mice. Together, these results suggest that an aged liver is in a state of “precarious balance”, due to concurrent activation of oncogenic and tumor suppressor pathways, but protected against neoplastic progression by IFN-signaling. Age-dependent activation of IFN signaling has been observed in many tissues and recent studies have demonstrated its detrimental consequences on aging, raising the question as to why IFN-signaling is activated during aging. We propose that aged tissues are intrinsically at higher risk of cancer and age-dependent activation of IFN-signaling is an adaptive process to protect from tumorigenesis, but one that also has maladaptive consequences.

## Introduction

Aging is a major risk factor for hepatocellular carcinoma (HCC). Incidence of HCC increases ∼75-fold from 30 to 70 years of age (*1*). However, the reasons for this striking age-dependence are not well understood. A better understanding of aging and its role in cancer can lead to improved risk assessment, early detection and prevention. This is a challenge to decipher because aging is a complex process whereby accumulation of damage, functional exhaustion and/or spontaneous “drift” of dynamic systems triggers adaptive changes that can be beneficial on one level but detrimental on another (*2, 3*). Deconstructing the relative impact of damage, adaptive and maladaptive processes on diseases of aging, including cancer, is a primary objective for the field. Previous studies have shown that age-dependent activation of interferon (IFN) signaling is, paradoxically, a cause of tissue degeneration (*4, 5*). This raises the critical question as to why this apparently maladaptive IFN signaling is commonly activated in aged tissues (*6–9*). Our data suggest that age-dependent activation of IFN-signaling/ISGs is an adaptive tumor suppression mechanism to counter the increased risk of tumorigenesis in aged tissues, associated with a precarious balance of dual activation of oncogenes and tumor suppressors.

## Results

To understand how aging affects mouse liver, we first examined gross physiological changes with age. Examination of Picro-Sirius red staining of healthy young (4-7 months of age) and old (22-25 months of age) mouse liver showed a higher level of collagen deposition, consistent with fibrosis, in aged liver (Fig 1A, B). Elevated liver stiffness, consistent with fibrosis, with age was also observed by Parallel Plate Shear Rheology (Fig 1C). H&E sections showed elevated levels of immune infiltration (Sup Fig 1-1A), confirmed through FACS analysis by a 2-3-fold increase of in Cd45+ cells, along with altered proportions of various immune cell types with age (Fig 1D, Sup Fig 1-1A-D). We observed a marked increase in Cd8+ T cells and monocytes, while at the same time a reduction in neutrophils, B cells, and Cd4+ T cells (Fig 1D-E). We further observed elevated populations of Pd1+ Cd8+ T cells and Tnfα+ Ifnγ+ Cd8+ T cells indicating an apparent activated population, along with Tox+, Tcf7-, Cd8+ T cells and Cd101+ Pd1+ Cd8+ T cells indicating simultaneous T cell exhaustion (Fig 1F-G, Sup Fig 1-2 A-D)(*10–13*). Metabolomic analysis of whole liver revealed a broad range metabolomic changes with age, including features of elevated phospholipid biosynthesis, spermidine biosynthesis, ubiquinone biosynthesis, and other metabolic pathways including methylhistidine, sphingolipid and glutathione metabolism (Fig 1H, Sup Fig 1-3 A, B, Sup Table 1). Epigenetic analysis of isolated hepatocytes from healthy aged mice revealed numerous differences compared to young, including DNA hypomethylation at enhancer regions confirming our previous findings in Cole et al., 2017 (*14*) (Fig 1I). Additionally, we identified the top 25 pathways linked to hypomethylated genes with age at CpGs within promoters and gene bodies as involved with numerous cancer-associated pathways, metabolism, mitophagy, bile secretion and cytokine signaling pathways (Fig 1J, Sup Fig 1-3 C-E). Chromatin immunoprecipitation of Histone H3K27ac showed the majority of genic regions, including exons, 3’UTR and intron, to be similar between age groups; however, there was a significant reduction of H3K27ac at promoter regions and an increase at intergenic regions in old mouse hepatocytes (Sup Fig 1-4 A). RNA seq analysis of whole liver identified large transcriptomic changes with age. The top transcriptional regulators predicted to be involved included Stat1, Stat3, Ctnnb1, Rela, Trp53, AP1 members (Fos and Jun), along with Myc (Fig 1K, Sup Fig 1-4B-C, Sup Table 2). Activation of Stat1 is consistent with activated IFN signaling (Sup Fig 1-4C). Most notably, these data suggest simultaneous activation of both oncogenic (e.g., Fos/Jun and Myc) and tumor suppressive (e.g., Trp53) pathways in the same aged liver, in the absence of evident neoplasia.

**Figure 1:**
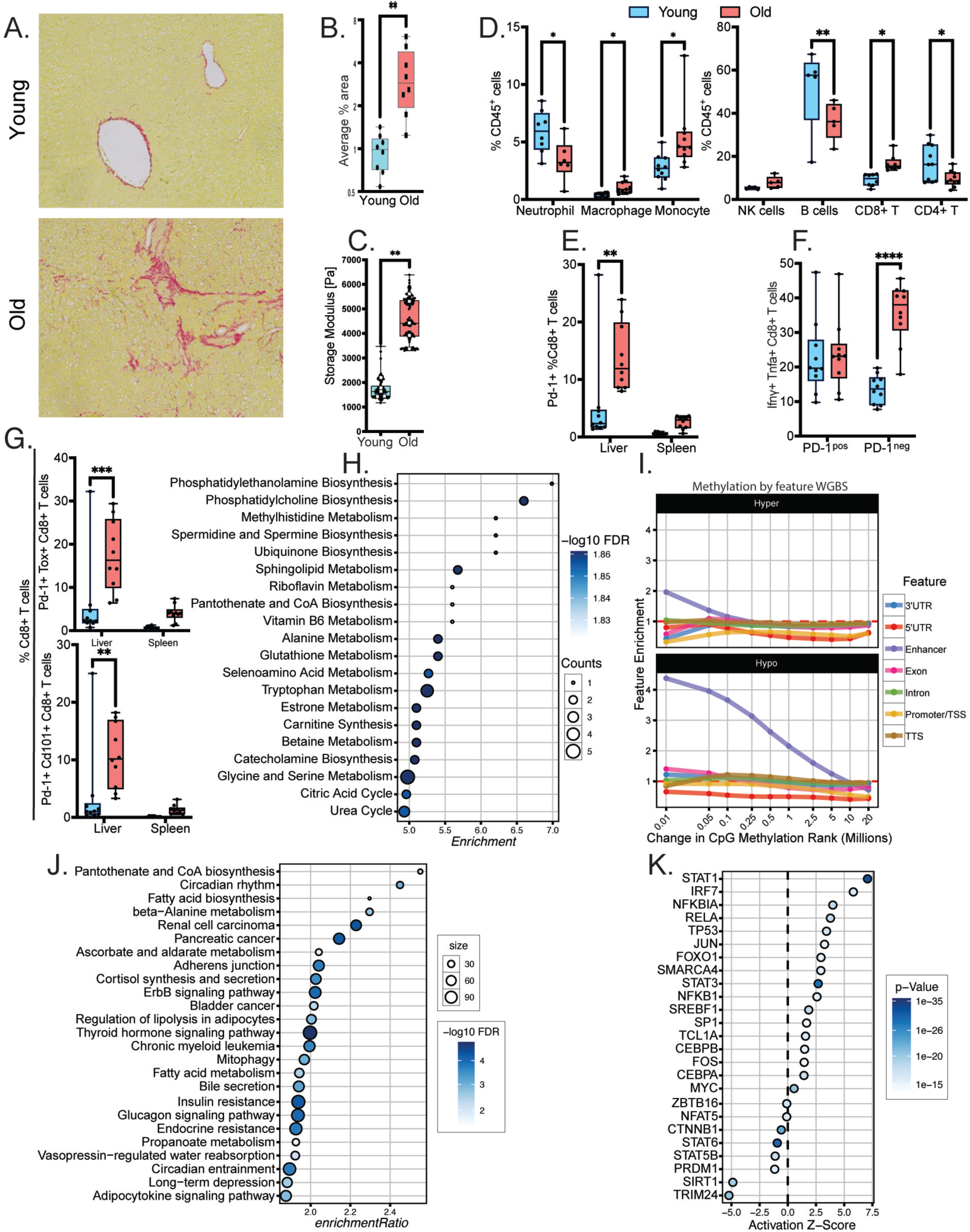
Aging changes the liver at the tissue and molecular level. A) Representative Picro-Sirus red images of young (7-MO, N=8) and old (25-MO, N=8) mouse liver showing elevated fibrosis with age. B) Picro-Sirus red staining quantitation. C) Parallel Plate Shear Rheology testing stiffness of young (2-MO, N=5) and old (22-MO, N=5) mouse livers. D) Box and violin chart of FACS immune profiling in young (5-MO N=5) and old (22-MO, N=5) mouse liver. E) % Pd-1+ Cd8+ T cells (activated). F) % lfny+ Tnfa+ Cd8+ T cells (activat­ ed) G) % Pd-1+ Cd101+ Cd8+ T cells and Pd-1+ Tox+ Cd8+ T cells (exhausted). H) Metabolomic KEGG pathway analysis comparing old (22-MO, N=10) and young (5-MO, N=9) mouse liver with indicated −log10 FDR and counts within pathway by size and color. I) Enrichment in CpGs rank-ordered by differential methylation significance between young (5-MO, N=4) and old (22-MO, N=4) isolated mouse hepatocytes, by genetic feature. J) KEGG pathways identi­ fied to be hypomethylated by lnfinium mouse methylation bead chip analysis comparing young (5-MO, N=5) and old (22-MO, N=5) isolated mouse hepatocytes. K) Ingenuity Pathway Analysis {IPA) of top 25 transcriptional upstream regulators predicted to be modulated in whole liver from old (25-MO, N=5) and young (7-MO, N=4). Statistical analysis represented using ANOVA in which asterisk (*) denotes p < 0.05, (**) p < 0.01, (***) p < 0.001 and (****) p < 0.0001.

**Figure 2:**
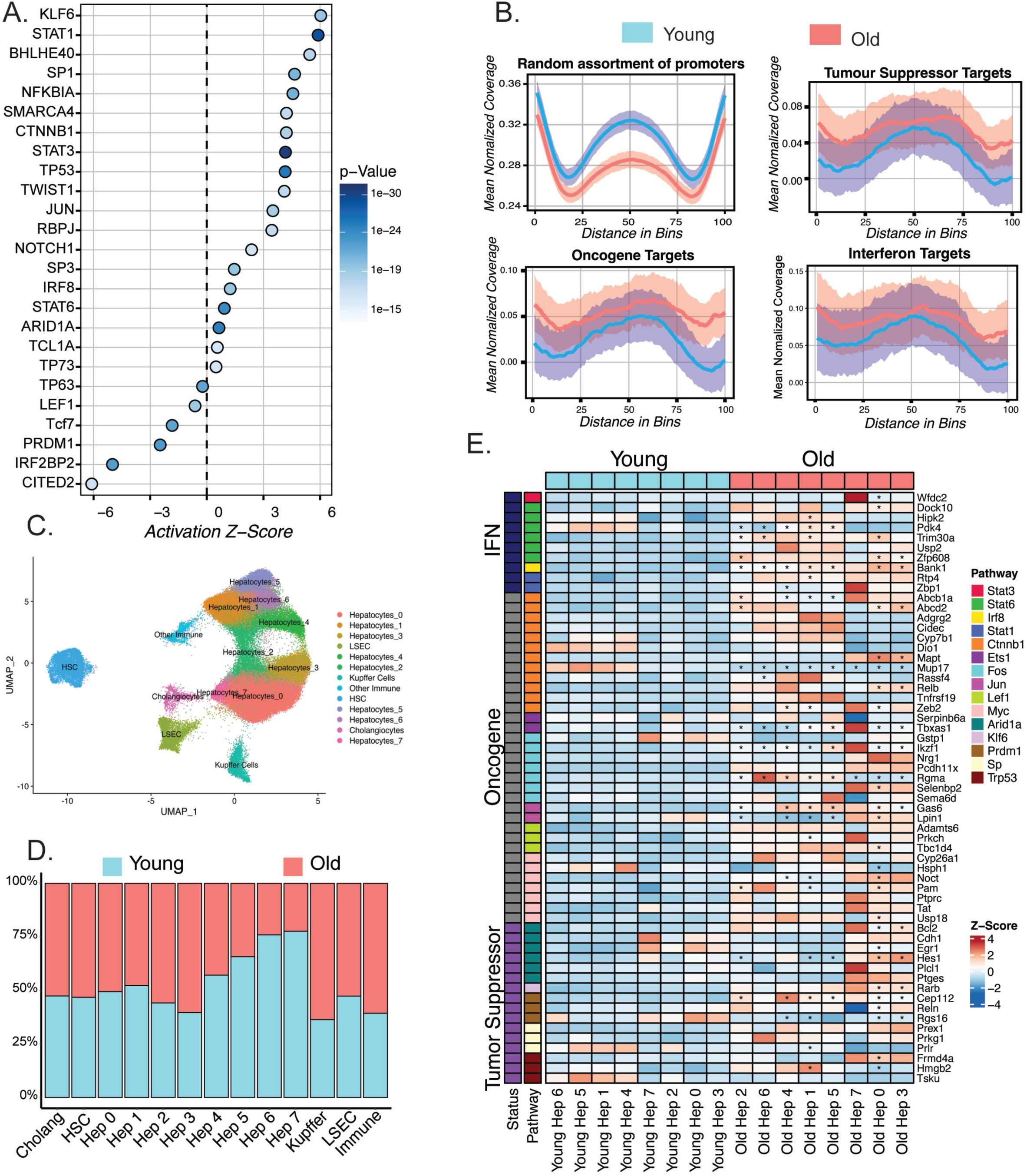
Aged liver shows dual activation of oncogenes and tumor suppressors in same cell clusters, suggestive of a "precarious balance". A) IPA of top 25 predicted upstream transcriptional regulators of pathways differentially expressed comparing old (22-MO, N=9) and young (5-MO, N=10) RNAseq of isolated hepatocytes. B) ChlPseq analysis of Histone H3K27ac deposition at promoters for random vs target genes of oncogenic and tumor suppressor pathways as predicted by IPA comparing old (22-MO, N=S) and young (5-MO, N=S) isolated hepatocytes. C) UMAP with indicated sub-clusters of snRNAseq comparing young (7-MO, N=4) and old (24-MO, N=4) mouse liver. D) Distribution of snRNAseq subclusters by age. E) Heatmap of oncogene and tumor suppressor genes elevated with age by hepatocyte subcluster in snRNAseq.

**Figure 3:**
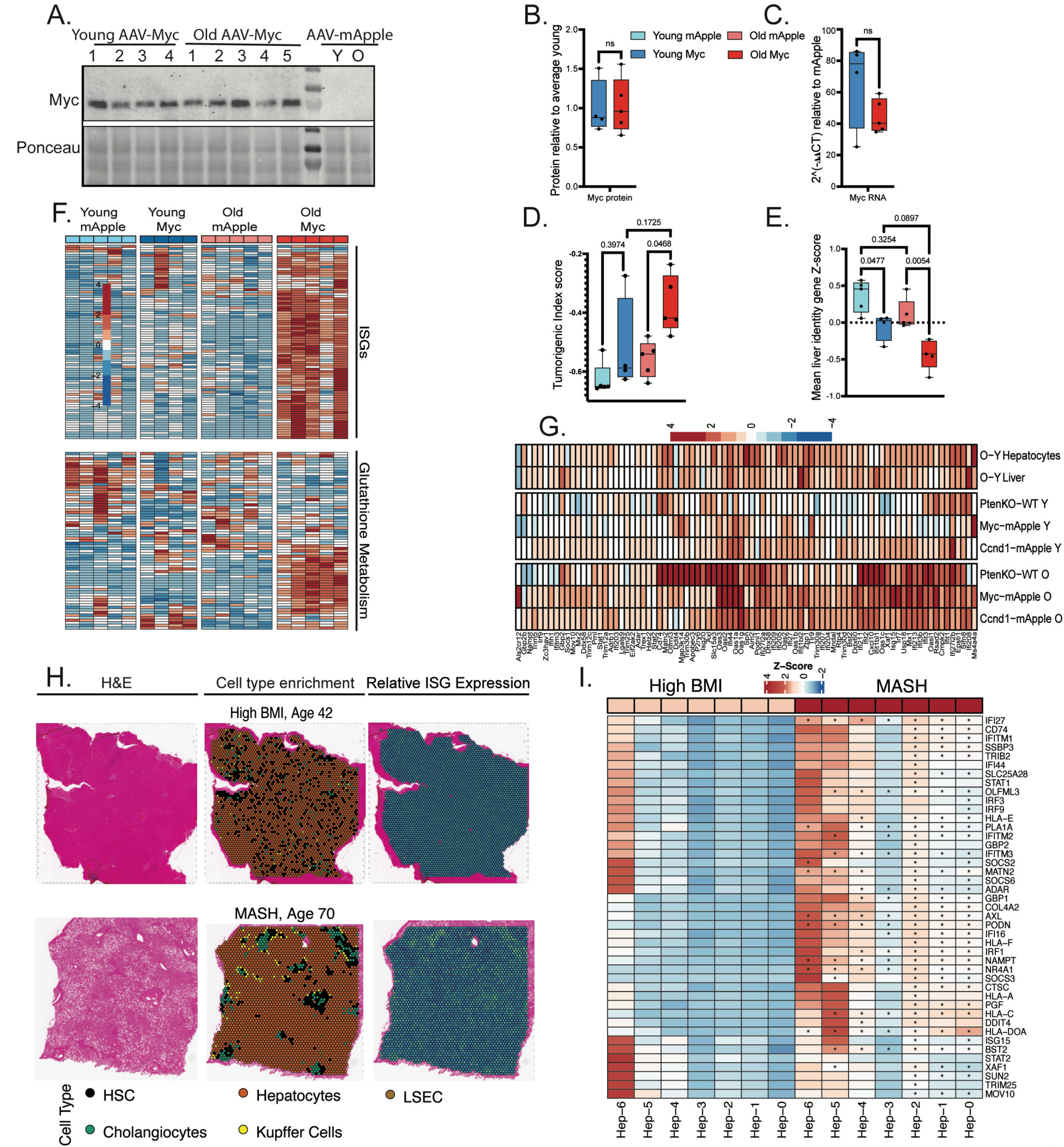
Oncogene activation in aged liver further elevates expression of ISGs. A) Representative western blot of Myc expression in liver of young (6-MO, N=4) and old (23-MO, N=5) mice infected for 30-days with indicated AAV. B) Quantitation of Myc western blot normalized to average young AAV-TBG-Myc infected. C) qPCR of AAV directed expression of Myc in young and old mice D) Calculated tumorigenic index by age and treatment. E) Liver identity gene expression score by age and treatment. F) Heatmap of ISGs and Glutathione metabolism gene expres­ sion by age and treatment. G) Comparative analysis of ISGs in young Ccnd1 (N=10) vs mApple (N=10), old Ccnd1 (N=8) vs mApple (N=9), Pten ko young (N=3) vs WT (N=3) and Pten ko old (N=9) vs age matched WT (N=3) mice. H) Represen­ tative Visium spatial images indicating H&E, cell type identification and ISG expression. I) Heatmap of ISGs as detected by Visium in different cell types showing log2FC comparing MASH (N=5) to high BMI (N=4) samples. Statistical analysis represented using one-way ANOVA with post hoc Tukey test in which asterisk(*) denotes p < 0.05, (**) p < 0.01.

**Figure 4:**
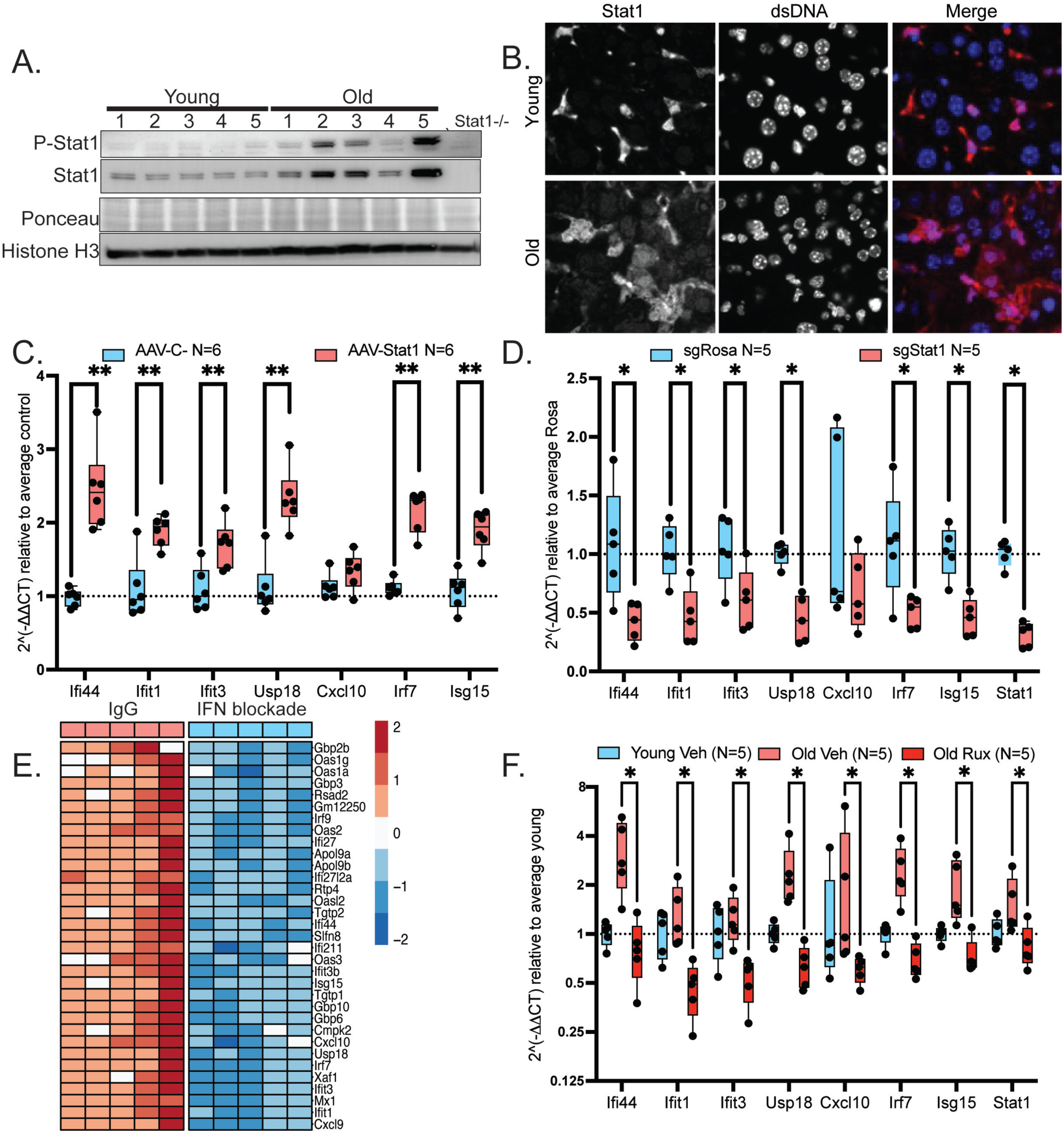
Expression of ISGs in aged liver is mediated by STAT1 and IFN signaling. A) Representative western blot of young (5-MO, N=5) and old (22-MO, N=5) hepatocytes for Phospho-Ser727 Stat1, total Stat1 with Ponceau Sand total Histone H3 as loading control. B) lmmunofluorescence of Stat1 expression in young (5-MO) and old (22-MO) mouse liver FFPE sections. C) qPCR of ISGs in young (6-MO) liver at 21-day post-infection of AAV-Stat1/3xFLAG (N=6) as compared to control AAV-MCS (N=6). D) qPCR of ISGs in old (25-MO) liver treated for 21-day with AAV-saCas9 directed knockout of sgRosa26 (N=5) control and sgStat1 (N=5). E) Heatmap of ISGs detected using RNAseq of old (24-MO) mice that received combination of anti-lrnaR1 and anti-lfn[g!R1 blockade (N=5) or isotype control blockade (N=5) for 21-days. F) qPCR of ISGs with 3-weeks ruxolitinib treatment in old (22-MO) mice as compared to vehicle old and vehicle young (5-MO, N=5/cohort). Statistical analysis represented using one-way ANOVA with post hoc Tukey test in which asterisk (*) denotes p < 0.05, (**) p < 0.01.

To test whether this phenomenon represents cell type heterogeneity in the liver or is a feature of hepatocytes, we first performed bulk transcriptome analysis of young and old purified mouse hepatocytes. The top transcriptionally activated pathways as indicated by Ingenuity Pathway Analysis (IPA) included upregulation of tumor suppressor pathways, such as Trp53, Klf6 and IFN signaling (Stat1, Stat6 and Irf8) (Fig 2A, Sup Fig 2-1A, Sup table 3). At the same time, we observed an elevation in oncogenic signaling, for example Stat3, Ctnnb1 and Jun pathway activation (*15–31*). We confirmed elevation of key oncogene and tumor suppressor targets in isolated hepatocytes from additional young and old mice using qPCR, specifically Fos, Jun, Ccnd1, p16, p53 targets (p21, Gadd45a, Tigar, Puma, Mdm2)) and ISGs (Ifi44, Ifit1, Ifit3, Usp18, Cxcl10, Irf7, Isg15, Stat1) (Sup 2-1B-F). ISGs were further confirmed to be elevated in mouse hepatocytes using Tabula Muris Senis(*32*) (Sup Fig 2-1G). Supporting this gene expression data, ChIP-seq analysis in isolated hepatocytes showed that, although most gene promoters appeared to lose H3K27ac with age, H3K27ac accumulated at most promoters of oncogenic and tumor suppressor target genes and IFN transcription factors (Fig 2B, Sup Fig 2-2A-C). To better resolve activation of oncogene and tumor suppressor pathways we performed single nuclei RNA-seq. Subcluster analysis showed that a majority of the hepatocyte subclusters increased expression of IFN signaling (Stat1, Irf7 and ISGs, e.g., Ifi44, Ifit3, Ifit1). Remarkably, most hepatocyte subclusters had simultaneous expression of both oncogenic and tumor suppressor genes and/or their target genes (Fig 2C-E, Sup Fig 2-3A). For example, oncogenic pathways Ccnd1, Ctnnb1 and Vegfa, and tumor suppressor p53 target genes (e.g., Mdm2 and Bbc3), indicating that in aged liver dual activation of oncogene and tumor suppressor pathways occurs in the same clusters of hepatocytes, suggestive of a “precarious balance” that is poised for transformation.

Given the seemingly poised state of the aged liver for transformation through dual activation of oncogenic and tumor suppressive pathways, we wondered whether aged liver might be more sensitive to oncogenic challenge as compared to young. To test this, we utilized Adeno-associated virus 2/8 (AAV) to ectopically express either mApple or c-Myc, the most frequently amplified oncogene in HCC (*15*), driven by a hepatocyte-specific thyroxine binding globulin (TBG) promoter. This induced expression in >99% hepatocytes (Sup Fig 3-1A)(*33–35*). Healthy young 4-month-old and old 22-month-old male wildtype C57BL6 mice were infected with retro-orbital injections of 1e^11^ virus particles/mouse. Mice were culled and livers were collected 30-days post infection. Gross analysis showed no overt neoplasia in young or old mice. Western blot and qPCR showed similar c-Myc expression in both young and old mouse liver (Fig 3A-C). Analysis of DNA methylation showed only a few Myc-induced alterations, with age being the strongest driver of changes in the methylation landscape (Sup Fig 3-1B). RNAseq analysis was conducted to determine age-specific differences in gene expression of c-Myc or Ccnd1 vs mApple. All groups were readily able to be distinguished from each other by PCA analysis (Sup Fig 3-1C). Old Myc showed the highest number of differentially expressed genes (DEGs) compared to any other group, with 259 up and 53 down regulated genes vs young Myc (Sup Fig 3-1D). Previously, *Wang et al.,* published an algorithm to determine “tumorigenic index (TI)” from RNAseq data, as a risk assessment for HCC in human and mouse tissues (*36*). Using this approach, we observed an increase in TI in both young and old with the expression of Myc, but only old Myc mice were significantly elevated as compared to their mApple age-matched control (Fig 3D). As cell transformation is typically accompanied by a loss of differentiation. Therefore, we assessed liver identity gene expression in our samples by comparing expression of liver-specific genes as previously defined by *Kim et al.,* and found Myc significantly reduced liver identity gene expression in both young and old mice with a stronger trend in old mice compared to young (Fig 3E, Sup Fig 3-1E)(*37*). We next wanted to know which pathways were specifically differentially regulated by Myc expression in old mice compared to young mice. String network analysis of RNA-seq data showed two major clusters of gene expression changes, specifically elevated ISGs and glutathione metabolism genes in c-Myc-infected old liver (Fig 3F, Sup Fig 3-1F-G). We observed similar elevation of the ISG signature in old mice challenged by another oncogenic stress, 30 days expression of Ccnd1 by AAV8-TBG-Ccnd1 (Fig 3G, Sup Fig 3-2A-C). In addition, we examined the ISG signature in an age-dependent model of Pten-/- tumorigenesis, and similarly observed an increase in ISGs dependent in Pten KO that was exacerbated by age (Fig 3G) (*36, 38*).

To assess the relevance of this ISG gene signature in human liver, we examined published RNAseq data, from Suppli et al., 2019, of liver from healthy, obese, MAFLD and MASH human donors. This showed an elevated ISG signature with disease progression (Sup Fig 3-3E)(*39*). To assess whether this ISG signature might also be linked to MAFLD progression in human liver, we examined Visium spatial RNAseq from patients classified by pathological analysis as “High BMI” and “MASH” with mean ages of 38.5 and 72.5 respectively (Sup Fig 3-3A). Cell types enriched within the tissue region were identified bioinformatically (Fig 3H, Sup Table 3) and gene expression was determined in hepatocytes, compared to the “High BMI cohort”. This analysis showed an elevation of oncogene and tumor suppressor gene and gene target expression linked to MAFLD progression and likely age (Sup Fig 3-3B-D). Chief among the signatures were ISGs which increased within the aged MASH cohort (Fig 3H, I). While we were unable to separate the effects of MASH and age within this tissue cohort, given the robust age gap between “high BMI” and “MASH”, age is likely a key biological factor for MASH progression. Taken together, these results show that in mice, aging substantially alters the response to increased oncogene activity, specifically resulting in a pronounced enhancement of ISG expression in the aged liver. This is paralleled in human liver where increasing risk of HCC reflected in MAFLD is accompanied by increased expression of ISGs, especially in the aged MASH tissues.

In light of the emphatic role of age in driving ISG expression, we set out to define requirements for ISG expression in aged liver. We focused on hepatocytes since they are the most abundant cell type in the liver and showed robust age-dependent expression of ISGs (Sup Fig 2-3B-C). To probe the role of the top predicted ISG regulating transcription factors, Stat3, Stat1 and Irf3, in aging hepatocytes, we profiled their protein expression in isolated hepatocytes of young and old mice. Neither Stat3 nor Irf3 protein level changed significantly in old hepatocytes as compared to young, but both total Stat1 and phosphorylated pS727 Stat1, indicating Jak activation, were elevated with age, as visualized by western blot of isolated hepatocytes (Fig 4A, Sup Fig 4-1A).

Immunofluorescence of Stat1 in whole liver tissue sections of young mice showed expression in smaller non-hepatocyte cells, but very little expression in hepatocytes (Fig 4B). However, old liver showed high cytoplasmic and nuclear expression of Stat1 in hepatocytes (identified by size and morphology), indicative of transcriptionally activated Stat1.

Next, we aimed to test whether Stat1 is necessary and sufficient for expression of ISGs in liver of wild type C57Bl6J mice. To test sufficiency, we utilized AAV2/8-TBG-Stat1/3xFLAG to express flag-tagged Stat1 in hepatocytes. We infected young mice with AAV-TBG-Stat1/3xFLAG, or AAV-TBG-MCS as control, and then collected whole liver tissues at 7, 14, 21, 60 and 120-days post infection. Western blot analysis confirmed expression of Stat1/3xFLAG protein for at least 120 days (Sup Fig 4-1B). qPCR analysis showed that Stat1 expression was elevated by day 7 post-infection and sustained to at least 120 days post infection (Sup Fig 4-1C). Expression of Stat1 resulted in upregulation of known ISG targets by day 14 to at least day 120 post AAV infection (Fig 4C, Sup Fig 4-1D for full time-course). To test whether Stat1 is required for ISG upregulation in old wild type liver, we knocked out Stat1 *in vivo* by infection of 25-month-old wild-type mice with AAV2/8-TBG-saCas9:U6-sgStat1 (or sgRosa26 as control). Mice were euthanized 3-weeks post AAV infection and whole liver collected. DNA analysis and western blot of Stat1 confirmed successful knockout (Sup Fig 4-1E, F). qPCR showed significant reduction of ISGs in Stat1 knock out vs control mice (Fig 4D). Additionally, we confirmed suppression of ISGs in old constitutive knock out genetically modified Stat1-/- mice (Sup Fig 4-2A). These results show that elevated Stat1 is both necessary and sufficient to upregulate ISGs in old wild type mouse hepatocytes.

To test whether expression of ISGs in old liver is ligand dependent, we treated old mice with combined anti-Ifnγ and anti-IfnaR1 antibodies or isotype control for 3-weeks. Mice were euthanized and RNA-seq was conducted. This showed that expression of ISGs was reduced in the Ifnγ/IfnaR1 blockade samples as compared to control (Fig 4E, Sup Fig 4-2B). Similarly, inhibition of Ifn receptor signaling by dosing with the Jak1/2 inhibitor, Ruxolitinib at 45 mg/kg, for three weeks also showed significant reductions in ISGs as compared to vehicle control (Fig 4F, Sup Fig 4-2C, D). Over-night treatment with ruxolitinib of isolated hepatocytes cultured e*x vivo* from young and old mice reduced ISGs (Sup Fig 4-2E), suggesting type I IFN signaling as hepatocytes do not express IFNγ. Confirming a role for type I Ifn signaling, we also observed a significant reduction of ISGs in old Ifnar1-/- mouse liver as compared to age-matched controls (Sup Fig 4-2F). Based on these results, we conclude that expression of ISGs in old liver is mediated by type I IFN signaling and STAT1.

Given the documented role of age-activated IFN signaling in driving diseases of aging, we wanted to understand the role of activated IFN signaling in an aged liver that appears poised for transformation(*40–42*). To test this, we repeated the experiment in young and old mice where we expressed either mApple or Myc while also providing oral gavage of ruxolitinib or vehicle 2x daily (5 days on+2 days off) for a 30-day span (Fig 5A). Western blot and qPCR confirmed upregulation of Myc at the RNA and protein level in the appropriate mice (Sup Fig 5-1A-B). qPCR analysis and RNAseq of liver from treated mice showed a reduction of ISGs in mice treated with ruxolitinib as compared to vehicle mice, confirming that ruxolitinib inhibited Jak/Stat signaling (Sup Fig 5-1B-C). We performed weighted gene co-expression network analysis (WGCNA) of the RNA-seq data which revealed disparate biological responses within the cohort of aged mice with Myc + ruxolitinib treatment. Specifically, this analysis identified two clusters with one group exhibiting a “tumor-like” transcriptional profile, characterized by elevated signatures of drug metabolism, chemical carcinogenesis, glutathione metabolism and xenobiotic metabolism, while the other “normal” group was more similar to old Myc + vehicle (Sup Fig 5-1D-E). Notably, the “tumor-like” Myc + ruxolitinib samples were more refractory to suppression of ISGs by ruxolitinib as compared to the “normal” Myc + ruxolitinib samples (Sup Fig 5-1F). Most importantly, the old “tumor-like” samples showed a relatively higher degree of similarity to signatures of both human HCC mined from TCGA and Pten KO mouse tumors (Fig 5B), confirming a tumor-like transcriptome which was only present in aged mice with both Myc and ruxolitinib treatment. This data indicates that age dictates a more tumor-like transcriptome in mice administered Myc and ruxolitinib.

**Figure 5:**
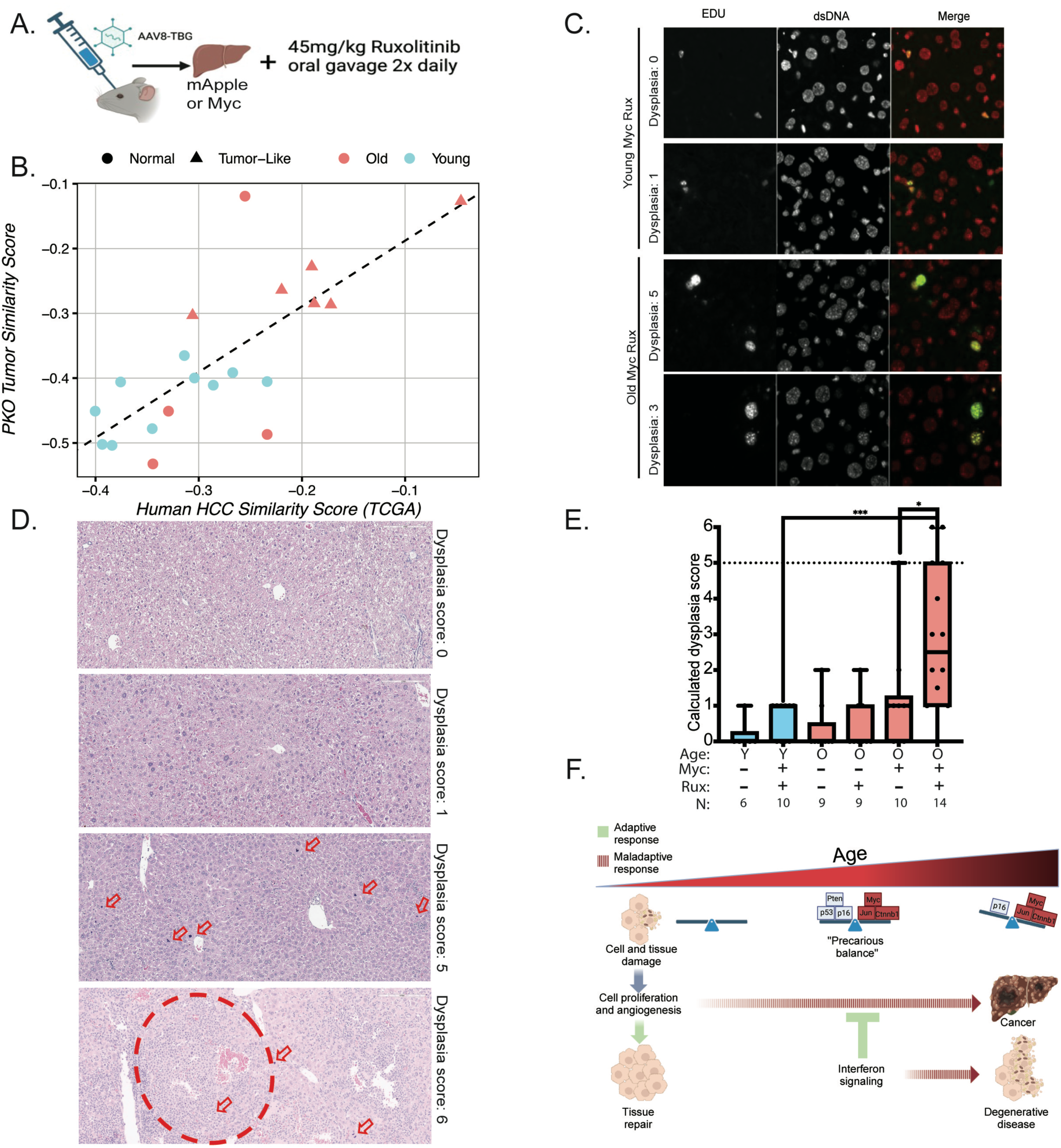
IFN signaling protects aged liver from tumorigenesis. A) Graphical representation of experimental design. B) Plot comparing median (Z(normalized counts)) of two gene sets for AAV-TBG-Myc+ruxolitinib treated mice. Set 1 (x-axis)= all genes upregulated in human HCC vs. normal (TCGA); Set 2 (y-axis)= all genes upregulated in mouse Pten KO tumors vs. wildtype.p-value for old vs young (N=10/cohort): Pten KO=0.0246; TCGA=0.0078. C) Representative images of EdU incorporation after 36-hours EdU exposure in young and old mice that received AAV-TBG-Myc and ruxolitinib. D) Representative H&E indicating regions of transformation (circle) and abnormal mitotic figures (red arrows). E) Graphical representation of blinded histopathological analysis of hepatocyte dysplasia and transformation by age and treatment. A dysplasia score >1 indicates elevating degrees of dysplasia and >5 is tumorigenesis. Each dot represents a mouse. F) Graphical representation of age-associated "precarious balance" and maladaptive response to tissue damage. Statistical analysis represented using one-way ANOVA with post hoc Tukey test in which asterisk (*) denotes p < 0.05, (**) p < 0.01, (***) p < 0.001 and (****) p < 0.0001.

EdU pulse labelling for 36-hours prior to analysis showed increased number of proliferating hepatocytes in old Myc + ruxolitinib tissues (Fig 5C, Sup Fig 5-2A). This increase is consistent with excess mitotic figures observed in hepatocytes in H&E of old Myc samples, with significantly higher levels in the old Myc + ruxolitinib as compared to old Myc + vehicle, indicating increased proliferation in these normally post-mitotic cells. Blinded histopathological analysis of these mouse livers revealed that 2 out of 10 Myc + vehicle old mice and 0 out of 10 Myc + ruxolitinib young had high grade dysplasia, but no tumors. In contrast, 10 out of 14 Myc + ruxolitinib old mice showed high grade hepatocyte dysplasia, with two harboring tumors (Fig 5D, E and Sup Fig 5-2B-G). H&E sections show representative images of treatment groups (Fig 5D), indicating regions of tumor, and examples of cellular abnormalities used to determine hepatocyte dysplasia including: anisokaryosis, anisocytosis, multinucleation, mitotic figure count and abnormal mitotic morphology. Immunofluorescence of tumor regions showed morphologically abnormal cells to be positive for markers of hepatocytes, including Hnf4a and Albumin, while negative for Cd45, confirming hepatocyte origin (Sup Fig 5-3). We conclude that, based on transcriptome and histological analysis, IFN inactivation *via* ruxolitinib treatment promotes tumorigenesis specifically in old mice expressing c-Myc and treated with Ruxolitinib. This shows that IFN signaling in aged mice serves as a tumor suppressive mechanism.

## Discussion

Our study reveals that aged hepatocytes exhibit an apparent “precarious balance” of dual activated oncogenic and tumor suppressor pathways, alongside increased IFN signaling. Young and old hepatocytes mount distinct responses to oncogenic stress, with aged hepatocytes showing exacerbated ISG expression upon oncogene activation. In humans, a similar ISG signature is observed in livers of aged donors with MASH. Stat1 is both necessary and sufficient for ISG expression in aged hepatocytes. Importantly, our findings indicate that activated IFN signaling serves a tumor-suppressive role in the aged liver, highlighting its protective function against tumorigenesis in aging.

Age-associated IFN upregulation has been observed in numerous aged tissues suggesting it to be a common feature of aging, albeit one whose function has not been established (*6–9*). Recent publications from Gulen et al. (Nature, 2023) describe elevated ISGs as sources of microglial inflammation, neuronal loss, cognitive impairment, and a driver of neurodegeneration in naturally aged mice (*4, 43*). Similar findings have been reported for elevated IFN signaling and aging phenotypes in the lung and intestine (*5, 41, 42, 44, 45*). These studies indicate that elevated ISG levels have detrimental effects on aging tissues and contribute to development of age-related pathologies. In contrast, our data suggest a beneficial role for upregulation of ISGs through suppression of tumor formation in the aged liver. Together, our data suggest that the apparent maladaptive age-associated activation of IFN signaling is likely an inadvertent consequence of adaptive activation of IFN signaling by age and oncogenic stress for the purpose of tumor suppression.

## Methods

### Animal usage

All animal procedures were approved by the Institutional Animal Care and Use Committee (IACUC) of Sanford Burnham Prebys Medical Discovery Institute. Animal experiments were performed at Sanford Burnham Prebys Medical Discovery Institute Animal Facility in compliance with the IACUC guidelines. The studies performed within this manuscript were performed with all relevant ethical regulations regarding animal research. Young and old C57BL/6J animals were obtained from the NIA aging colony housed at Charles Rivers. Additional young C57BL/6N mice were purchased from Charles Rivers. Animals were housed 5 mice per cage and maintained under controlled temperature (22.5°C) and illumination (12h dark/light cycle) conditions. Teklad Gloabal 18% Protein Rodent diet (Envigo, ref 2018) was provided ad libitum. Stat1-/- as described by Durbin et al., Cell 1996 from our internal breeding colony were used in studies described (*46*).

Ifnar1-/- (Jackson Laboratories, Strain 028288) mice were housed according to the University of California at San Diego IACUC facilities. Mice were maintained in 12h light/12h dark cycle, ambient temperature of 20–22°C, relative humidity of between 30 and 70%, with unrestricted access to water and fed a standard chow diet (Lab Diets 5001) ad libitum.

### Mice for liver stiffness

C57BL/6J female mice were acquired at 4 weeks of age from Jackson labs (Strain: 000664) and housed according to the University of California at Berkeley Animal Care and Use Committee standards in housing facilities. Mice were maintained in 12h light/12h dark cycle, ambient temperature of 20– 22°C, relative humidity of between 30 and 70%, with unrestricted access to water and fed a standard chow diet (Lab Diets 5001) ad libitum.

### Human liver tissues

were obtained through the Biorepository and Laboratory Services (BLS) at the Clinical and Translational Science Institute at University of Minnesota (IRB#00013764). Upon collection by the surgeon, the liver specimens were placed in 10% formalin and further processed with standard FFPE processing procedure at BLS core. The hematoxylin and eosin (H&E) and trichrome staining sections from each FFPE block were reviewed and graded by a hepatic pathologist.

### IFN blockade

was provided to 22-month-old mice for 21 days at 10mg/kg IP at a total volume of 100μl alternating sides every Monday, Wednesday and Friday (total of 9 injections). Antibodies were purchased from Leinco clones and resuspended in sterile PBS treatment group 1:anti-IFNγ (XMG1.2), anti-IFNAR1 (MAR1-5A3) or treatment group 2: anti-mouse IgG2b (HKSP) and anti-rat IgG1 (G113). Mice were separated into treatment groups at random and all administrations were conducted at 1pm.

### Ruxolitinib treatment

Ruxolitinib was purchased from LC Laboratories (R-6600) provided twice daily oral gavage dissolved in vehicle consisting of 2% DMSO, 30% PEG300 and 68% sterile H2O. Mice were weighed every Monday, Wednesday and Friday to during administration period to insure health and safety of treatment. Mice were separated into treatment groups at random and all oral gavage administrations were conducted 7am and 3pm daily with a holiday on Saturdays and Sundays (*47*).

### Plasmids

pAAV[Exp]-TBG>mApple:WPRE, AAV-TBG-MCS_108bp, pAAV[Exp]-TBG>mCcnd1[NM_001379248.1](ns)/3xFLAG:P2A:mApple:WPRE, pAAV[Exp]-TBG>3xFLAG/mMyc[NM_010849.4](ns)*:P2A:mApple:WPRE and pAAV[Exp]-TBG>mStat1[NM_001205313.1]*/3xFLAG:WPRE were generated from VectorBuilder. pX602-AAV-TBG::NLS-SaCas9-NLS-HA-OLLAS-bGHpA;U6::BsaI-sgRNA originated from Addgene plasmid #61593.

### SaCas9 sgRNA design and validation

Guide RNAs were generated using CRISPick with mouse reference genome GRCm38 (Ensembl v.102). sgRNA were cloned into pX602-AAV-TBG::NLS-SaCas9-NLS-HA-OLLAS-bGHpA;U6::BsaI-sgRNA and transformed into NEB Stable competent E. Coli. Insertion was validated via direct colony (Genewiz) using their universal U6 primer 5’-GACTATCATATGCTTACCGT-3’. Knockout was confirmed in mouse genomic DNA extracted using Monarch Genomic DNA Purification kit (New England Biolabs, Ref T3010S) followed by PCR amplification of the surrounding ∼500bp next to the sgRNA target site. PCR was performed using 2X Taq PCR Master Mix (APExBIO, Ref K1034), 100ng of gDNA and 1μl of 100uM primers. PCR products were then purified using QIAquick PCR Purification Kit (QIAGEN, Ref 28104) and sequenced via sanger sequencing with the forward amplification primers (Genewiz). To quantify cut efficiency, ICE analysis was performed as described in *“Conant, David et al.* 2019 (*48*). v3.0. Synthego; [04/15/2024].

### AAV generation and usage

Large-scale rAAV productions were performed by the SBP Functional Genomics Core Facility. In short, HEK-293T cells were co-transfected with rAAV vector, RepCap 8, and Helper (Cell Biolabs) plasmid DNA *via* polyethylenimine (Polysciences). After 96 hours post-transfection, crude viruses were isolated from supernatant through polyethylene glycol precipitation and cell pellet *via* sonication, followed by benzonase (Millipore Sigma) treatment. Concentrated viral particles were purified by iodixanol (Millipore Sigma) gradient ultracentrifugation. The fraction containing rAAV particles was collected and buffer-exchanged with PBS before being aliquoted and stored at −80°C. The viral titer was determined by SYBR green QPCR.

### Hepatocyte isolation

was conducted similar to described in Charni-Natan et al., 2020 (*49*). Mice were euthanized with CO2 asphyxiation. Once immobilized, mice were opened and a 24Gx3/4” (Surflo SR-0X2419CA) catheter was inserted into the inferior vena cava below the liver. 50ml of HBSS (Gibco Ref 14175-095, pre-heated to 42°C) at a flow-rate of 6ml/min then digested with 40ml DMEM (Gibco Ref 10313-021) containing Collagenase Type IV (Gibco, Ref 17104-019) pre-heated 42°C. livers were excised then disassociated in ice-cold DMEM with 2% Bovine Serum Albumin (BioWorld Cas: 9048-46-8) and stored on ice. Hepatocytes were washed using centrifugation at 50xg for 2 min and media changed at 4°C 5x then resuspended in 40% Percoll (Cytiva Ref 17089102), 10% 10x HBSS (Gibco Ref 14065-056) and 60% DMEM then centrifuged for 7 min at 100xg at 4°C. Viable pellet at the bottom of the centrifuge tube were collected, washed 1x in DMEM at 50xg then quantitated using hemocytometer with trypan blue stain.

### Ex vivo hepatocyte Ruxolitinib treatment

hepatocytes were isolated (as discussed previously) and cultured on type I Collagen (Gibco, RefA10483-01) coated plates and maintained at 37°C, 5% CO2 incubator. Hepatocytes were initially plated in media containing 10% FBS in DMEM with Pen-Strep. After hepatocytes were adhered to plastic overnight, the media was removed and replaced with DMEM with Pen-Strep without FBS containing ruxolitinib. Samples were collected with scraping directly in trizol or lysis buffer for western blot.

### Tissue homogenization

whole liver was homogenized either fresh from mouse or snap frozen using Precellys *Western blot:* homogenized tissue or cell suspension was quantitated using Bradford reagent (Pierce Ref 1863028) and Spectra Max 190 plate reader. Fresh SDS-PAGE gels were cast and as described previously (*50*). Using the BioRad mini Protean Tetra system and transferred to PVDF (Immobilion PSQ, ISEQ00010) for 70 min at 100V. Ponceau S stain (Sigma-Aldrich P7170-1L) was taken prior to blocking in 5% milk (BD, Difco skim milk, Ref 232100) TBST for 1 hour at room temperature. Primary antibodies (Suplimentary table 6) were incubated in 5% BSA (BioWorld, Ref 9048-46-8) overnight 4°C on rocker. The following secondary antibodies were used: Goat anti-Mouse IgG-HRP (Thermo Fisher Scientific 362 Cat#31446, RRID:AB_228318), Goat anti-Rabbit IgG-HRP (Millipore Cat#AP307P, RRID:AB_92641). Ponceau S and HRP imaging conducted using BioRad ChemiDoc touch imaging system with Image lab software for analysis and image production.

### Immunofluorescence

Tissue was fixated for 48 hours at room temperature with 10% Neutral Buffered Formalin (Epredia, Ref 9400-1). After fixation, liver was paraffin embedded and sectioned into 5um sections. Sections were deparaffinized with 2x washes in 100 xylene, 95% Ethanol, 90% ethanol and 70% ethanol followed by antigen retrevial using 10mM Tris, 1mM EDTA and 0.05% Tween-20 for 40 minutes in steamer (Nesco, ST-25F) followed by 15 min permibilization in 0.05% triton-x PBS and 2 hours of blocking in 4% BSA, 1% FBS at room temperature. Primary antibodies (Supplimental table 6) were applied over-night 4°C. secondary antibodies (supplemental table 6) were applied 1 hour at room-temp followed by 1 hour in 0.1% Sudan black (Sigma, 199664) in 70% ethanol for background subtraction. DAPI was applied at 5μg/ml for 15 min. Coverslips were mounted in Flouromount-G (ThermoFisher, Ref 00-4958-02) and imaged using Nikon T2 microscope.

### Histology and stains

*Picro-Sirus Red Stain* conducted on 5μm FFPE sections as described https://med.emory.edu/departments/medicine/divisions/cardiology/research/labs/microscopy-in-medicine/_documents/picrosirius-red-stain-collagen.pdf. EdU kit used as described by Salic et al., PNAS 2008 (*51*). EdU was provided 140μl/mouse in PBS inter-peritoneally 36-hours prior to euthanasia.

### Metabolomics

Mice were euthanized according to IACUC protocol. Immediately after euthanasia, the liver was weighed (nearest 0.1mg) frozen tissue specimens using ice-cold 5:3:2 methanol:acetonitrile:water to a final concentration of 15mg tissue per ml(*52*). Mixtures were vortexed 30 min at 4**°**C and centrifuged for 10 min at 18,000xg, 4**°**C. Samples were randomized then 10μL of extracts were injected into a Thermo Vanquish UHPLC system (San Jose, CA, USA) and resolved on a Kinetex C18 column (150 × 2.1 mm, 1.7 μm, Phenomenex, Torrance, CA, USA) at 450μL/min through a 5 min gradient from 0 to 100% B (mobile phases: A = 95% water, 5% acetonitrile, 1 mM ammonium acetate; B = 95% acetonitrile, 5% water, 1 mM ammonium acetate) in negative ion mode (*53*). Solvent gradient: 0-0.5 min 0% B, 0.5-1.1 min 0-100% B, 1.1-2.75 min hold at 100% B, 2.75-3 min 100-0% B, 3-5 min hold at 0% B. Injections were then repeated for positive ion mode at 450μL/min through a 5 min gradient from 5 to 95% B (mobile phases: A = water, 0.1% formic acid; B = acetonitrile, 0.1% formic acid) in positive ion mode. Solvent gradient: 0-0.5 min 5% B, 0.5-1.1 min 5-95% B, 1.1-2.75 min hold at 95% B, 2.75-3 min 95-5% B, 3-5 min hold at 5% B(*53*). Eluant was introduced to a Thermo Orbitrap Exploris 120 mass spectrometer using electrospray ionization. For both negative and positive polarities, signals were recorded at a resolution of 70,000 over a scan range of 65-900 m/z. The maximum injection time was 200ms, microscans 2, automatic gain control (AGC) 3 x 10^6 ions, source voltage 4.0 kV, capillary temperature 320**°**C, and sheath gas 45, auxiliary gas 15, and sweep gas 0 (all nitrogen). Resulting .raw files were converted to .mzXML format using RawConverter. Metabolites were assigned and peak areas integrated using El-Maven (Elucidata) alongside the KEGG database and an in-house standard library of >600 compounds (*52, 54*).

### qPCR

RNA was isolated using trizol (Invitrogen, Ref 15596018) according to the manufacturer’s recommendations with chloroform extraction. RNA was quantitated using Nanodrop one (Thermo Scientific) then converted to cDNA (RevertAid Reverse Transcriptase, Thermo Fisher Ref EP0441), Ribolock RNase Inhibitor (Thermo Fisher Scientific Ref EO0381), 5x reaction buffer for RT (Thermo Fisher) and gene expression quantified by a standard SYBR-based approach using PowerUp SYBR Green Master Mix for qPCR (Applied Biosystems Ref A25741) QuanStudio 6 Felx Real-Time PCR System, 384-well (Applied Biosystems Ref 4485691). Primers used for qPCR are included in supplemental table 7.

### Parallel plate Rheology

Viscoelastic properties of tissues were determined using an oscillatory rheometer (Anton-Paar Ref M302e) with parallel-plate geometry of 8mm and a gap height of ∼0.2mm under 0.1% strain and 1 Hz frequency at 37°C in a humidity-controlled chamber.

### Flowcytometry

Tissues were dissected and placed on ice-cold RPMI1640 supplemented with 10% FBS. Single-cell suspensions of immune cells from the liver were obtained by mechanical disaggregation through a 70µm cell strainer (VWR) and washed through with 10% FBS in RPMI. Liver samples were spun at 60 r.c.f. and 4°C for 2 min with no brake to pellet hepatocytes before Percoll (Cytiva) centrifugation. The supernatant was collected, spun at 420 r.c.f. and 4°C for 4 min. The pellet was resuspended with 40% Percoll (Cytiva) in HBSS to further remove debris and hepatocytes. The isolated immune cells from the liver went through red blood cell lysis with ACK buffer (KD Medical) before counting cells on a hematocytometer. Splenocytes were isolated by passing cells through a 70µm cell strained followed by red blood cell lysis with ACK buffer before being transferred to a 96-well U-bottom plate and resuspended in fluorescence-activated cell sorting (FACS) buffer (2% FBS in 1X PBS). Viability staining was performed using LIVE/DEAD fixable red stain (1 in 1000 in FACS buffer, Invitrogen) for 15 min at room temperature. Suspensions were then pelleted and resuspended in anti-CD16/32 antibodies (1:500, BioLegend) to block non-specific binding of Fc receptors. Cells were incubated with the indicated surface antibodies for 30 min at 4°C. A FoxP3 transcription factor staining kit (eBioscience) was used for intracellular staining. Antibodies against intracellular proteins were diluted in 1X permeabilization buffer and added for 45 min at 4°C.

### Cytokine staining

cells were stimulated with PMA (final concentration of 1µg/ml) and ionomycin (Iono, Cell Signaling; final concentration of 1µg/ml) for 4h at 37°C in the presence of brefeldin A (GolgiPlug, BD Biosciences; final concentration of 1µg/ml) to block cytokine export from the golgi apparatus. 2% paraformaldehyde (PFA) was used to fix the cells after staining. Cells were resuspended in 100µL 1X PBS and run on the LSRII flow cytometer (BD Biosciences). Data were analyzed using FlowJo (v.10, BD Biosciences).

### Pathological analysis

H&E whole slide images were provided to a board-certified pathologist who remained blinded to groupings. The pathologist identified neoplastic transformation in some tissues, and in subneoplastic tissues identified varying degrees of the following criteria of hepatocellular dysplasia: anisokaryosis, multinucleation with 3 or more nuclei, bizarre nuclei, increased mitotic rate, bizarre mitotic figures. Based on the range of abnormalities identified, a 0-6 dysplasia scoring scale was established with 0 indicating no elements of dysplasia, scores from 1-5 indicating the cumulative dysplastic criteria, and 6 indicating the presence of neoplasia.

### WGBS

Primary hepatocytes were isolated and snap frozen using liquid nitrogen, as described above, from 5-month-old and 22-month-old C57BL/6 mice. DNA was extracted using the Zymo quick-DNA miniprep kit (Zymo D3024) following manufacturer instruction. After DNA integrity was verified by tapestation it was then processed by BGI according to SOP-SS-023 version A0 in which bisulfite treatment followed DNA shearing, end repair, A-tail addition and methylated adapter ligation was conducted. PCR amplified libraries were sequenced using Novaseq system and down-stream analysis was conducted according to below.

*Infinium mouse BeadChip* (*46*) as directed by Illumina-cite manufacturer (Ref 20041558) on primary hepatocytes isolated and snap frozen using liquid nitrogen, from 5-month-old and 22-month-old C57BL/6 mice. DNA was isolated using Zymo quick-DNA miniprep kit (Zymo D3024) and submitted to UCSD IGM genomics core for downstream tapestation quantitation and processing according to manufacturer instructions and assayed using Illumina iScan.

### PolyA RNAseq

RNA was isolated using trizol (Invitrogen, Ref 15596018) according to the manufacturer’s recommendations with chloroform extraction. RNA was quantitated using Nanodrop one (Thermo Scientific) and tapestation. Library preparations were performed with the Watchmaker Genomics mRNA Library Prep Kit (Watchmaker Genomics) and Elevate Long UDI Adapters (Element Biosciences). RNA libraries were pooled, and sequenced (2×75bp) on the Element Biosciences AVITI sequencer with the 2×75 Cloudbreak kit. Or NEB polyA mRNA library preparation kits were used for library prep and sequencing with Illumina NextSeq500.

### ChIPseq

was conducted on isolated hepatocytes from C57BL/6J male mice that were snap frozen following protocol as described by Erb et al., Nature 2017(*55, 56*). In short, hepatocytes were lysed and chromatin sheared to between 250-500bp as confirmed by agarose gel electrophoresis in size using sonication (Biorupter pico, Ref B01060010). Protein G Dynabeads (Invitrogen, Ref 1000D) were prepared with 5μg of primary antibody of interest: Histone H3 (Abcam ref ab1791), Histone H3K27ac (Abcam, ref ab4729), Histone H3K4me1 (Abcam, ref ab8895). 5μg chromatin as quantitated by qubit dsDNA HS Kit (Invitrogen Ref Q32854) were used per ChIP and IP was allowed overnight 4°C while inverting. After washing steps, DNA was eluted using phenol chloroform isoamyl alcohol (Thermo Scientific, Ref J62336) isolation and ethanol precipitation then quantitated using qubit dsDNA HS Kit (Invitrogen Ref Q32854) and submitted to Sanford Burnham Prebys genomic core for library preparation and sequencing on NextSeq500.

### 10x single nuclei RNAseq

Single-nucleus HT RNA-seq was performed by the Center for Epigenomics at UC San Diego. Briefly, for each liver, a sample of tissue was homogenized using gentleMACS™ M tubes and gentleMACS™ Tissue Octo Dissociator. Samples were homogenized in dissociation buffer (5mM CaCl_2_, 2mM EDTA, 3mM Mg(CH_3_COO)_2_, 10mM Tris – HCl pH = 8.0, 0.6mM DTT, 1U/μl RNAse Inhibitor (N2515, Promega), 1X Roche cOMPLETE Protease Inhibitor, EDTA-Free (11873580001, Sigma)) using Protein 01_01 program. Nuclei suspension was incubated for 10 min at 4°C and filtered with 30μm filter (CellTrics). Nuclei were pelleted with a swinging bucket centrifuge (500xg, 5 min, 4°C; 5920R, Eppendorf) and resuspended in 500µL of sort buffer (1% Fatty acid free BSA (7500804, Proliant; 21-040-CV, Corning) in PBS, 1U/μl RNAse Inhibitor, 1X Protease Inhibitor) and stained with 7-AAD (1:100) for 20min on ice, protected from light. After staining nuclei were sorted using a SH800 (Sony) flow cytometer into 5X collection buffer (5% Fatty acid free BSA in PBS, 5U/μl RNAse Inhibitor). Next, nuclei were pelleted (500xg, 5 min, 4°), resuspended in 1X Nuclei Buffer (1X Nuclei Buffer (10X Genomics, 2000207), 1mM DTT, 1U/μl RNAse Inhibitor), and counted using a hemocytometer. About 33,300 nuclei were loaded onto a Chromium Controller (10x Genomics). Libraries were generated using the Chromium Single-Cell HT 3′ Library Construction Kit v3.1 (10x Genomics, 1000350, 1000351, 1000352) with the Chromium Single-Cell M Chip Kit (10x Genomics, 1000349) and the Chromium i7 Dual Index Kit TT Set A 96 rxns Kit for sample indexing (10x Genomics, 1000215) according to manufacturer specifications. Final library concentration was assessed by Qubit dsDNA HS Assay Kit (Thermo-Fischer Scientific) and fragment size was checked using TapeStation High Sensitivity D1000 (Agilent) to ensure that fragment sizes were distributed normally about 500 bp. Libraries were sequenced using the NextSeq500 and a NovaSeq6000 (Illumina) with the following read lengths: 28 + 8 + 91 (Read1 + Index1 + Read2).

### Visium Spatial transcriptomics

studies were carried out using two Visium platforms. For direct mount workflows, human liver sections were sectioned with 5μM thickness and directly mounted onto Visium slide (10x Genomics PN-2000233) and processed following 10x Genomics Protocol CG000409. For CytAssist workflow, human liver blocks were sectioned with 5μM thickness and mounted on standard frosted glass slides and processed following 10x Genomics Protocol CG000520. Slides were kept in desiccator within 1-5 days after sectioning until staining and processing. Briefly, slides were baked at 60°C for 2 hours before deparaffinization and rehydration, followed with H&E staining and slides were imaged on a Huron TissueScope LE scanner. In the Visium CytAssist workflow, after probe hybridization, the tissue sections mounted on the standard glass slides were aligned on top of the capture areas of the Visium slide (PN-1000519) placed in the CytAssist instrument. A brightfield image was captured to provide spatial orientation for data analysis, followed by hybridization of transcriptomic probes to the Visium slide. Probe release, extension and the remaining steps followed the standard Visium user guide for FFPE (Protocol CG000407 for direct mount; Protocol CG000495 for CytAssist). Library construction was generated following the manufacturer’s instructions as listed above for probe based FFPE library construction. In short, after probe ligation, products were captured and probes were extended by the addition of UMI, Spatially Barcode and partial Read1. The spatially barcoded probe products were released from the slide and harvested for qPCR to determine sample Index PCR cycle number, followed with indexing via Sample Index PCR. The final library molecules were cleaned up by SPRIselect and quantified by bioanalyzer. Libraries were sequenced on Illumina sequencing instrument NovaSeq 6000 or NextSeq 2000 according to 10x Genomics Visium manufacturer’s guidance, with targeted sequencing depth of minimum 25,000 read pairs per tissue covered spot on capture area. Standard Illumina binary base call (BCL) sequencing data output files were generated and converted to Fastq format.

### NGS analysis

#### Bulk RNAseq Analysis

For all experiments, raw reads were quality-checked with FastQC v0.11.8. Experiments containing adaptor sequences were trimmed with Trim Galore v0.4.4_dev. Reads were aligned to the mm10 reference genome with STAR aligner v2.5.3a (*57*) and converted to gene counts with HOMER’s analyzeRepeats.pl script (*58*). Gene counts were normalized and queried for differential expression using DESeq2 v1.30.0 (*59*). For each pairwise comparison, genes with fewer than 10 total raw counts across all samples were discarded prior to normalization, and genes with an absolute log2foldchange greater than 1 and an FDR-corrected p-value less than or equal to 0.05 were pulled as significant.

#### Functional Enrichment

Gene sets were queried for treatment-specific functional enrichment using over-representation analysis (ORA) and gene set enrichment analysis (GSEA) in WebGestaltR v0.4.4(*60*).

#### ChIPseq Analysis

For all experiments, raw reads were quality-checked with FastQC v0.11.8. Experiments containing adaptor sequences were trimmed with Trim Galore v0.4.4. As each sample was spiked in with human DNA, reads were aligned to a custom, combined mm10 and hg38 genome with STAR aligner v2.5.3a(*57*). Reads that aligned to the hg38 portion of the combined genome were filtered out of each sample. Duplicate reads were removed from each sample using Picard v2.18.4. Peaks for each file were calculated using HOMER’s findpeaks.pl script relative to each sample’s Input file(*58*). Metagenes were produced using metagene2 v1.12.0. ChIPseq coverage was normalized with the log2_ratio normalization method.

#### Whole Genome Bisulfite Sequencing Analysis

Raw reads were aligned with Bismark v0.15.0(*61*) and CpG Dyads were collapsed using the methods described in Cole et al, 2017(*14*). Differentially methylated regions were identified using methylKit v1.10.0(*62*), wherein significantly differentially methylated CpGs between conditions were called with a mean methylation difference above 25% and an FDR-corrected p-value below 0.001.

#### Metabolomics

Metabolomics data were processed using Metaboanalyst v5.0(*63*). Raw data were log10 transformed and auto-scaled (mean-centered and divided by the standard deviation). Metabolites with a raw p value less than 0.05 were designated as significantly different. Metabolites were included for quantitative enrichment analysis using the KEGG human metabolic pathways database (*64*). Pathways with an FDR less than 0.05 were considered significant.

#### snRNAseq

For each sample, shallow and deep sequencing fastq files processed using Cell Ranger v7.1.0(*65*). Filtered matrix files were processed using Seurat v4.3.0.1(*66*). Cells with nFeature less than 200, nFeature greater than 2500, or percent mitochondrial DNA greater than 1% were removed from each sample. Samples were normalized using SCTransform (*67*) and integrated following Seurat’s default pipeline using Harmony v1.2.0 (*68*). Cell types were evaluated using known canonical markers for cell types in the liver and using the markers for each cluster, as calculated by the FindMarkers function in Seurat. Each cluster as determined by Seurat (resolution = 0.8), was subclustered (resolution = 0.5) to identify clusters of nuclei that are likely not true nuclei. Any subcluster whose cell type could not be determined using known markers or markers calculated by Seurat were removed. Upon annotation, genes in each cell type were evaluated for differential expression between old and young. Genes with an absolute log2FC greater than 0.25 and an adjusted p value (Bonferroni correction) less than 0.05 were considered differentially expressed.

#### Visium

All samples were processed using Seurat v4.9.9.9.058(*69*). Spots with 0 were removed prior to analysis. In addition, any spot that was located two rows of spots from the edge of the tissue, including empty space within the tissue slice, were removed to reduce additional noise. Samples were normalized with SCTransform (*67*) and integrated following Seurat’s default pipeline using Harmony v1.2.0 for Visium data (*68*). Because the Visium spots have a diameter of 55μm, many spots likely have more than one cell type under them (*70*). Clusters were assigned the most likely cell type using canonical markers and markers calculated for each cluster. Upon annotation of spots, genes were evaluated for differential expression for each comparison. Genes were considered significant if the absolute log2FC was greater than 0.25 and the adjusted p value (Bonferroni correction) was less than 0.05.

#### Methylation

All methylation array samples were processed using SeSAMe v1.18.4 (*71*). CpG probes that are only in one group were removed prior to analysis. Loci were considered differentially methylated if their adjusted p value (Bonferroni-Hochberg correction) was less than 0.05.

