## Supplemental Figures for "Activated Interferon Signaling Suppresses Age-Dependent Liver Cancer"

A.

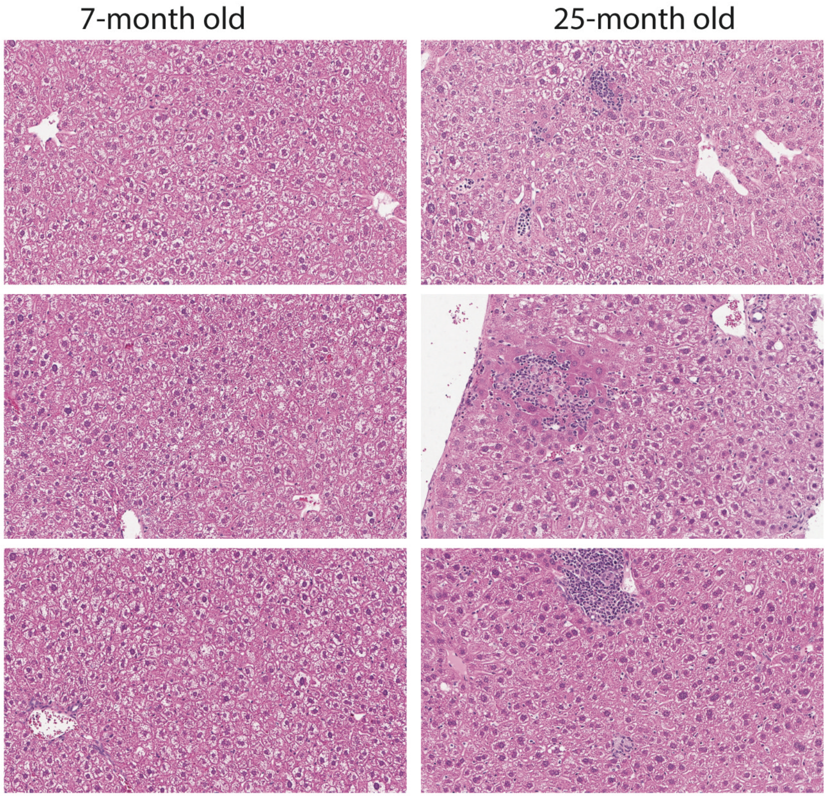


B.

| Age of mouse | Total cell number of CD45+ immune cells |
|---|---|
| 5 months | 3 million |
| 5 months | 5 million |
| 5 months | 3 million |
| **22 months** | **21 million** |
| **22 months** | **15 million** |
| **22 months** | **25 million** |

C.

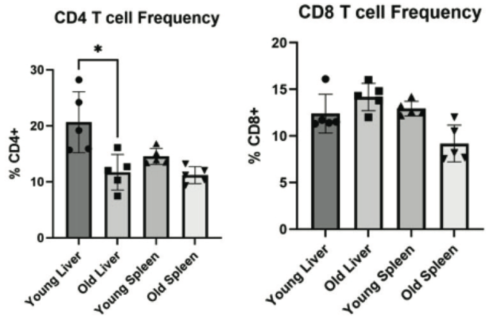


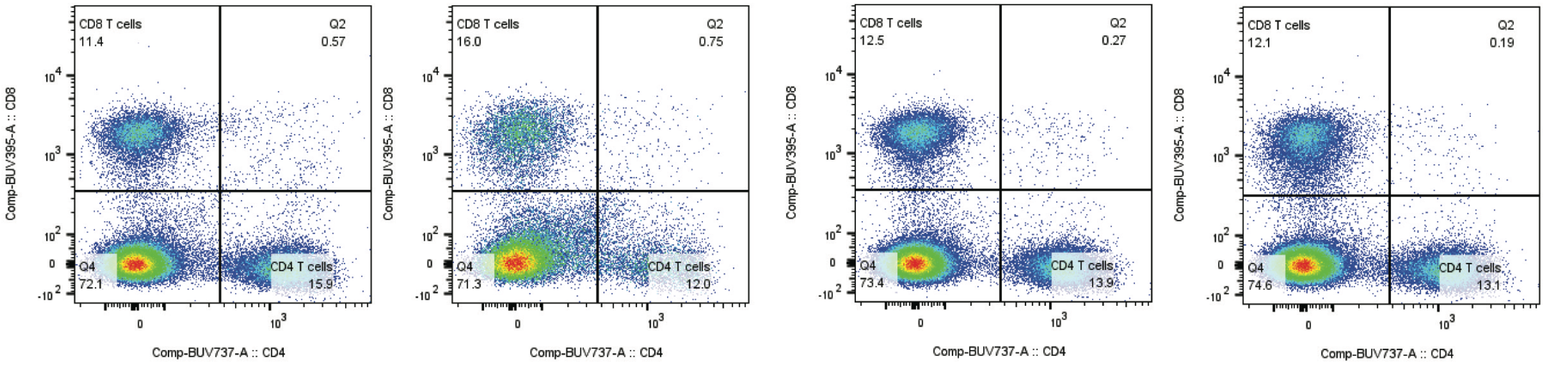


D.

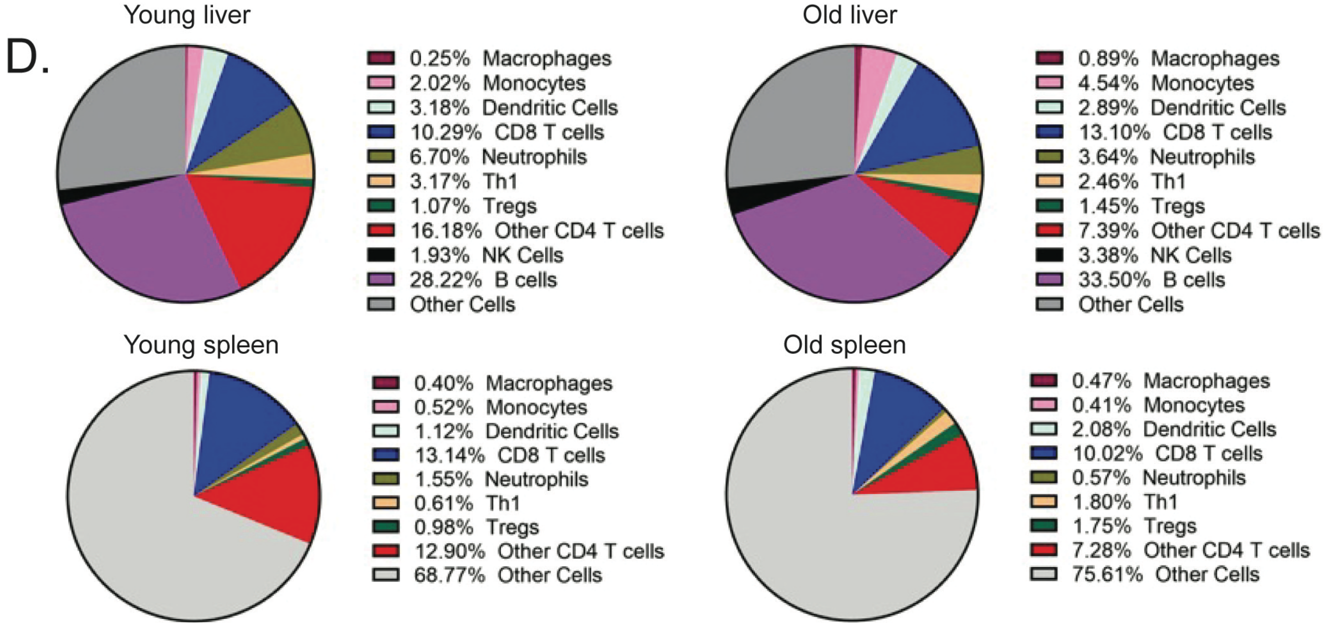


**Sup Fig 1-1: Age-associated elevated immune infiltration in liver.**
A) Representative images of H&E healthy young (7-MO, N=3) and old (25-MO, N=3) liver showing increased immune infiltration. B) Table showing quantitation of total events of Cd45+ cells from an equal portion of liver by weight comparing young (5-MO, N=3) and old (22-MO, N=3). C) Cd4+ and Cd8+ frequencies comparing young (5-MO) and old (22-MO) mouse liver as compared to spleen with representative FACS scatter plot below. D) Pie chart of percent Cd45+ cells comparing young (5-MO) and old (22-MO) liver. Statistical analysis represented using one-way ANOVA in which asterisk (*) denotes $p < 0.05$.

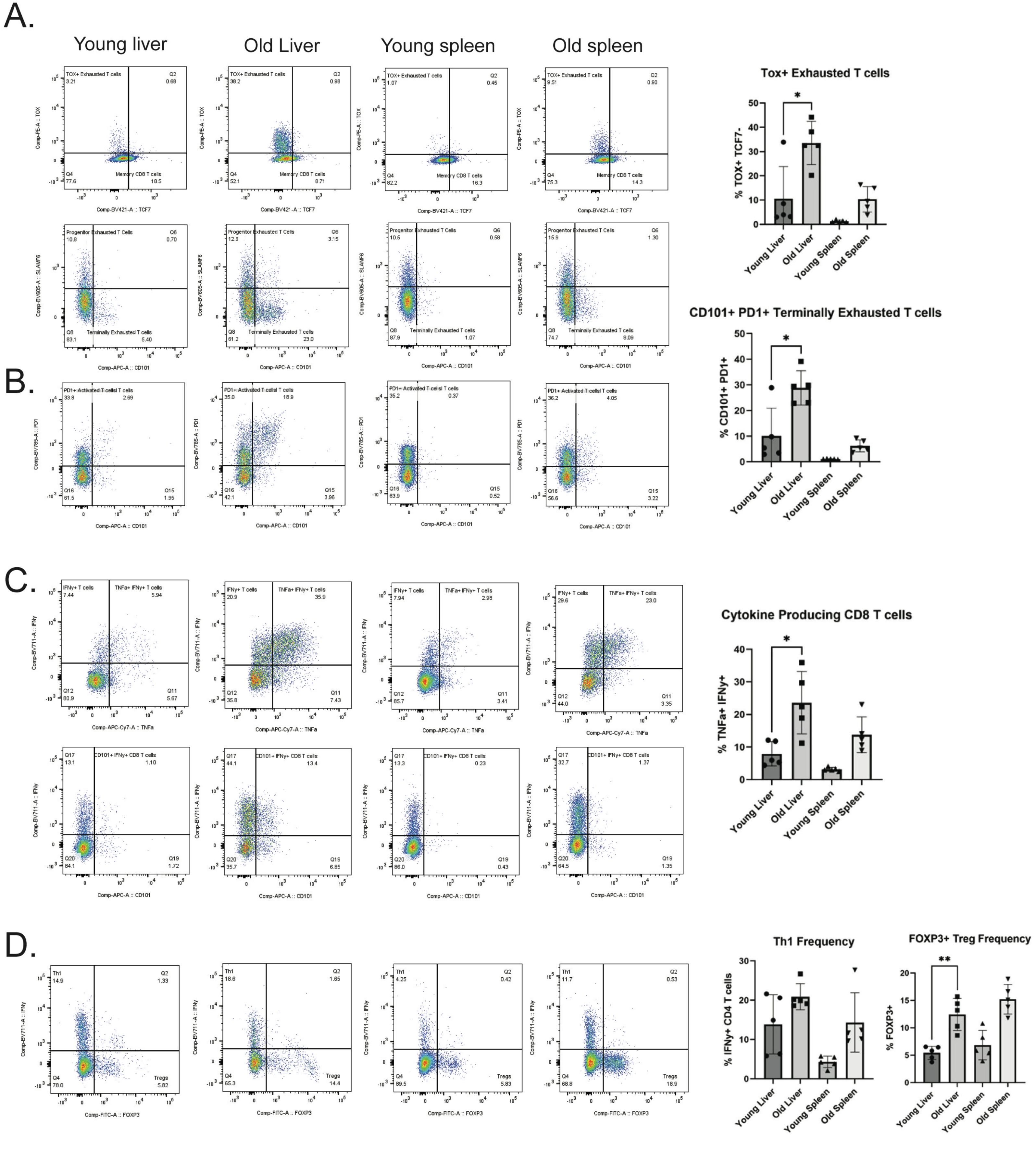


**Sup Fig 1-2: Increased markers of T-cell exhaustion and regulatory T-cells in aged liver.**
Representative FACS analysis comparing young (5-MO) and old (22-MO) with quantitation's for A) Tox+, Tcf7- Cd8+ T cell frequency, B) Cd101+ Pd1+ Cd8+ T cell frequency, C) Tnf⍰+, Ifn⍰+ Cd8+ T cell frequency, D) Ifny+ Cd4+ Th1 frequency and Foxp3+ Treg cell frequency in liver and spleen. Statistical analysis represented using one-way ANOVA in which asterisk (*) denotes $p < 0.05$.

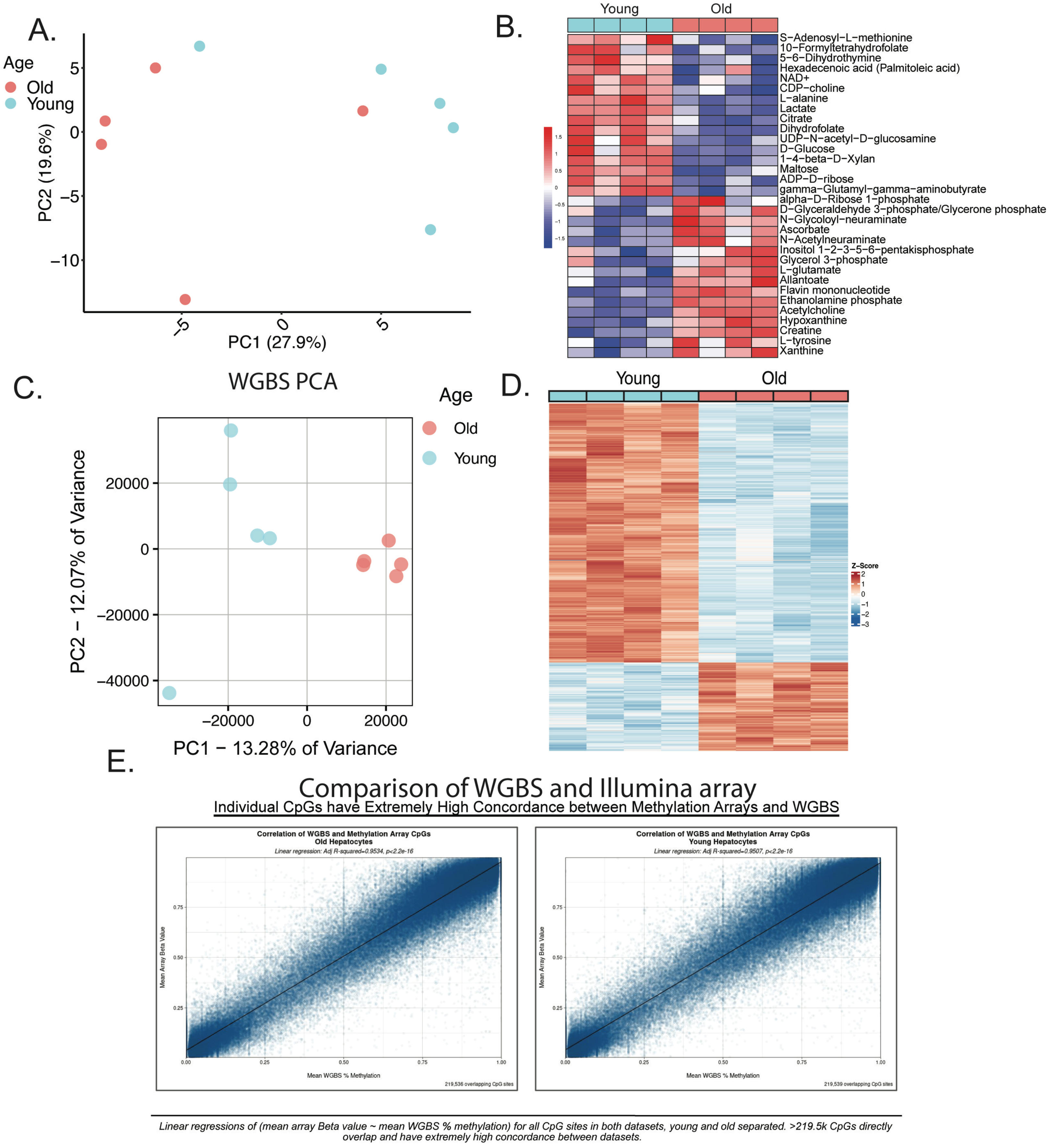


**Sup Fig 1-3: Age-associated metabolic and DNA methylation changes in mouse liver.**
A) Principal component analysis comparing metabolomic signature of young (5-MO, N=5) and old (22-MO, N=5). B) Heatmap of differentially detected metabolites by age. C) Principal component analysis comparing young (5-MO, N=4) and old (22-MO, N=4) isolated mouse hepatocytes differentially methylated loci by WGBS and D) heatmap of differentially methylated loci as detected by WGBS. E) Comparison of concordance between WGBS (% methylation) vs Infinium mouse methylation BeadChip (beta value) in which linear regressions are compared between data sets with an adjusted R2 value >0.95 and p>2.2e-16.

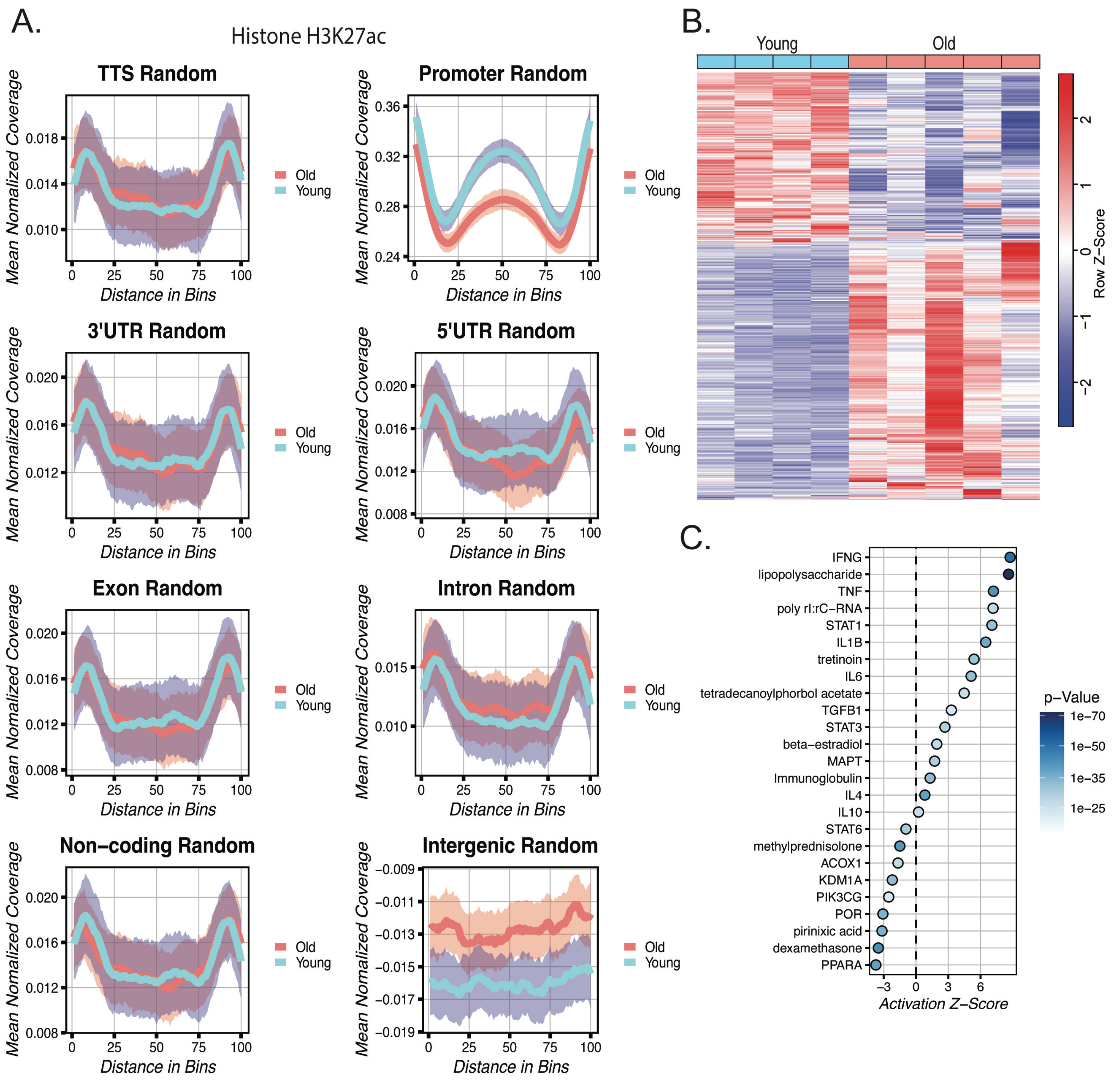


**Sup Fig 1-4: Enrichment of loss of Histone H3K27ac at promoters and altered transcriptome in mouse hepatocytes and liver.**
A) Representative genomic feature map comparing Histone H3K27ac ChIP for young (5-MO, N=5) and old (22-MO, N=5) isolated mouse hepatocytes. B) Heatmap comparing RNAseq of young (5-MO, N=10) and old (22-MO, N=9) isolated mouse hepatocytes. C) Ingenuity pathway analysis of predicted top 25 upstream regulators (unfiltered) detected from RNAseq with indicated activation Z-score and P-value by color.

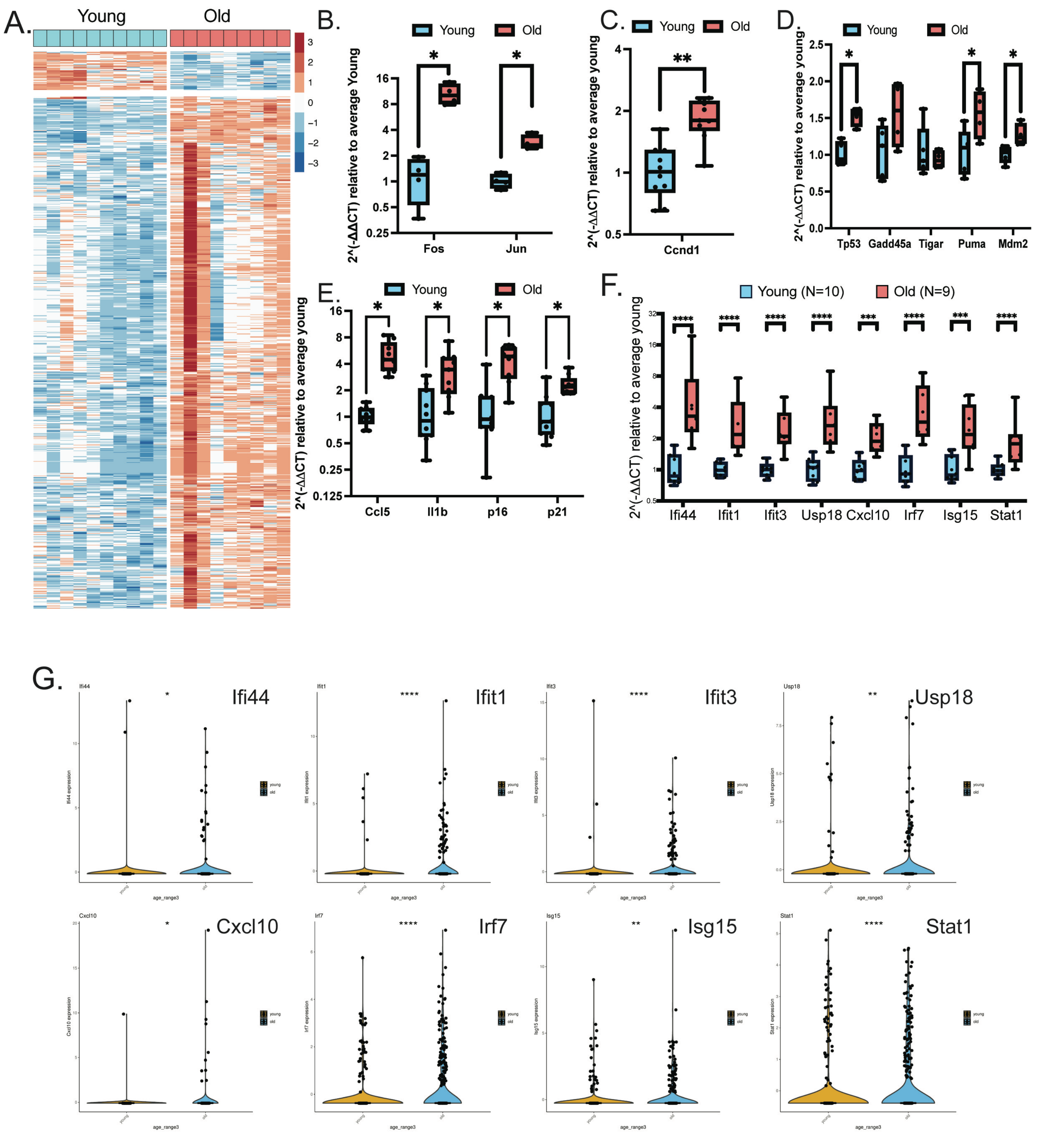


**Sup Fig 2-1: Altered expression of tumor suppressor, oncogene and IFN stimulated gene expression in hepatocytes with age.**
A) Heatmap of differentially expressed genes found in RNAseq of isolated hepatocytes from old (22-MO, N=10) and young (5-MO, N=9). qPCR measuring gene expression of an alternate set of hepatocytes isolated from old (22-MO, N=9) and young (5-MO, N=10): B) Jun, Fos. C) Ccnd1. D) p53 target genes, E) senescence associated secretory phenotype (SASP) F) IFN stimulated gene (ISG) expression in which each dot represents a different mouse. Statistical analysis represented using Mann-Whitney test in which asterisk (*) denotes $p < 0.05$, (**) $p < 0.01$. G) Plots representing ISG expression in Tabula muris senis. Statistical analysis represented using one way ANOVA with post hoc Tukey test in which asterisk (*) denotes $p < 0.05$, (**) $p < 0.01$.

A.

### Oncogene

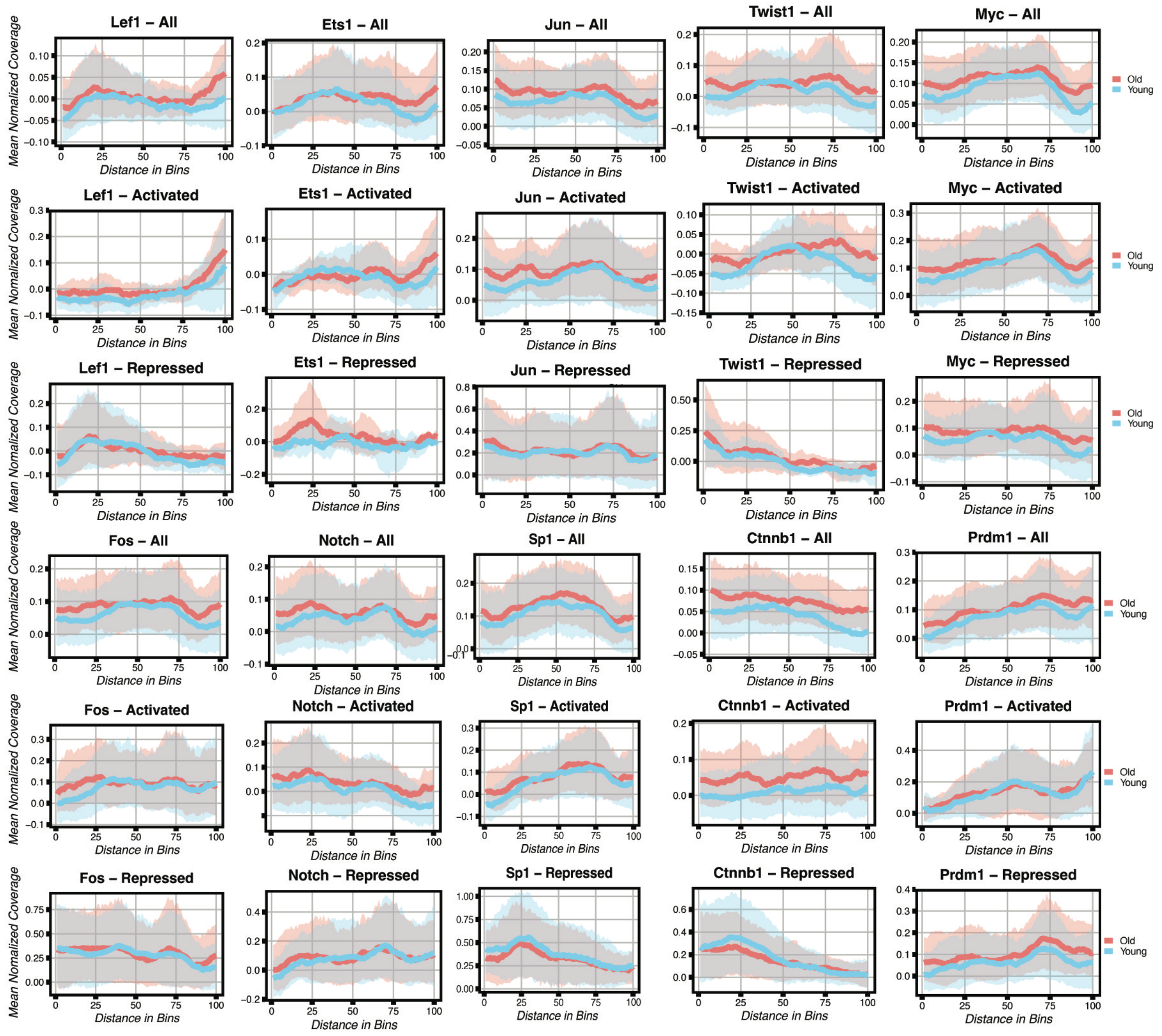


B.

### Tumor suppressor

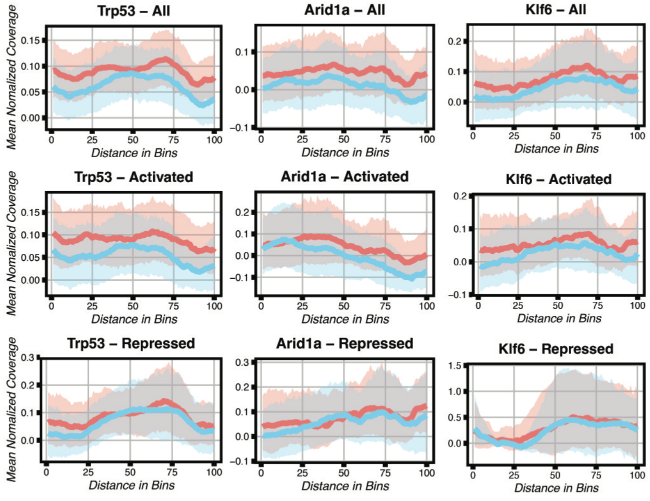


C.

### Interferon

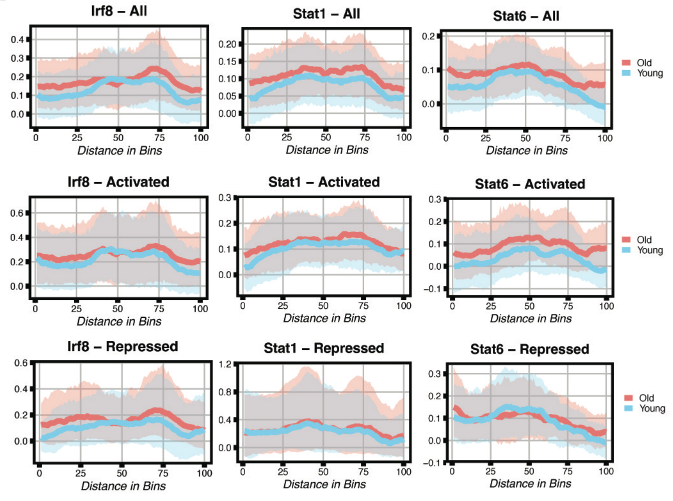


**Sup Fig 2-2: The distribution of Histone H3K27ac at oncogene, tumor suppressor and interferon target gene promoters do not decrease with age in hepatocytes.**
Histone H3K27ac deposition in old (22-MO, N=5) and young (5-MO, N=5) isolated hepatocytes at promoters of target genes for predicted A) oncogene target genes, B) tumor suppressor target genes and C) interferon target genes. Target genes as predicted by IPA are listed in supplemental table 5.

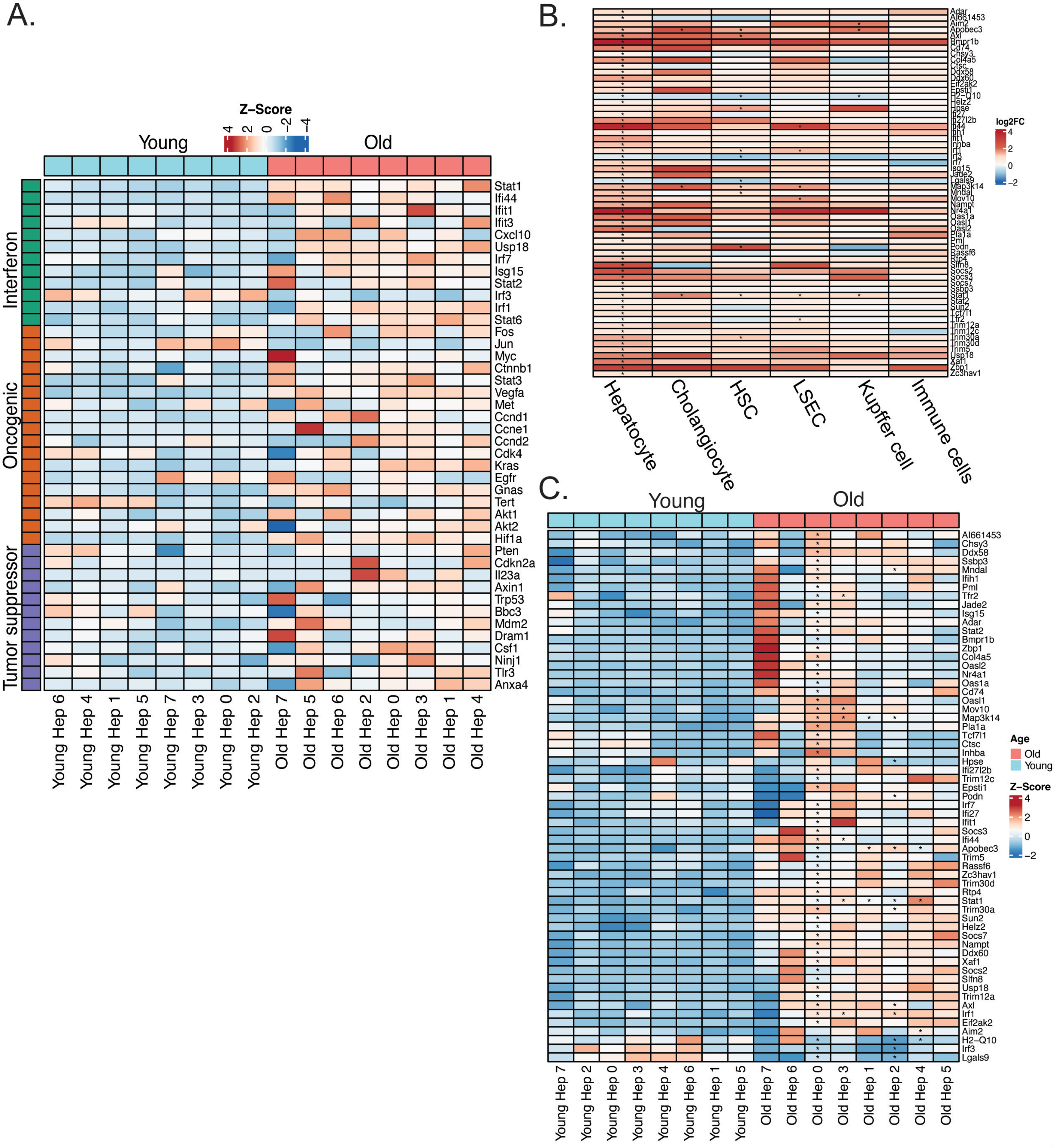


**Sup Fig 2-3: Elevated interferon, oncogene and tumor suppressor gene and target gene expression in old hepatocyte clusters.**

A) Heat map of interferon, oncogene and tumor suppressor gene expression using snRNAseq in old (25-MO, N=4) relative to young (7-MO, N=4) mouse hepatocyte clusters as predicted by IPA listed in supplemental table 5. B) ISG log2FC expression as compared to young for each indicated cell type. C) ISG expression in hepatocyte clusters as identified using snRNAseq in young and old. Statistical analysis represented using one-way ANOVA with post hoc Tukey test in which asterisk (*) denotes $p < 0.05$, (**) $p < 0.01$.

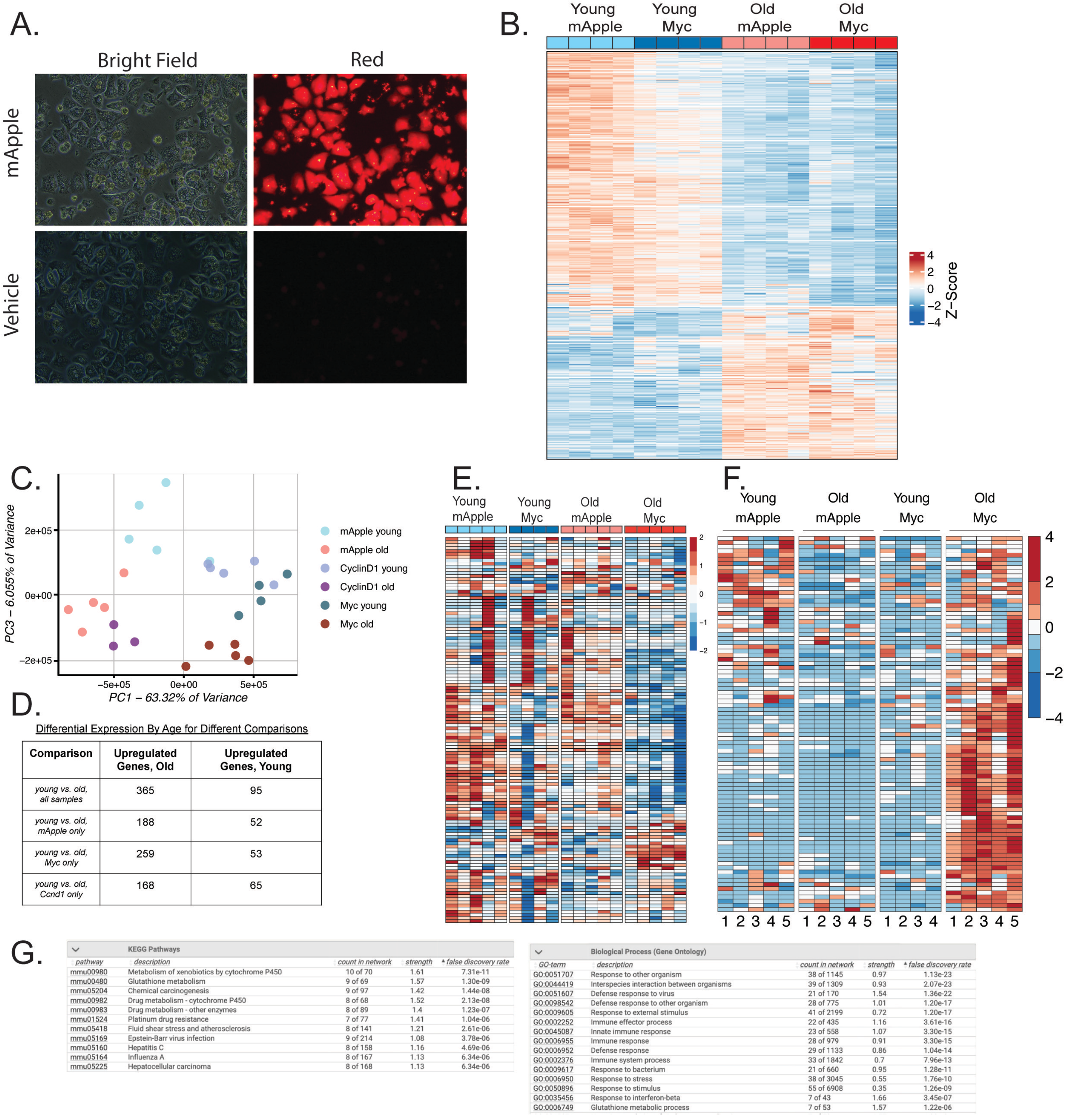


Differential Expression By Age for Different Comparisons

| Comparison | Upregulated Genes, Old | Upregulated Genes, Young |
|---|---|---|
| *young vs. old, all samples* | 365 | 95 |
| *young vs. old, mApple only* | 188 | 52 |
| *young vs. old, Myc only* | 259 | 53 |
| *young vs. old, Ccnd1 only* | 168 | 65 |

KEGG Pathways

| pathway | description | count in network | strength | false discovery rate |
|---|---|---|---|---|
| mmu00980 | Metabolism of xenobiotics by cytochrome P450 | 10 of 70 | 1.61 | 7.31e-11 |
| mmu00480 | Glutathione metabolism | 9 of 69 | 1.57 | 1.30e-09 |
| mmu05204 | Chemical carcinogenesis | 9 of 97 | 1.42 | 1.44e-08 |
| mmu00982 | Drug metabolism - cytochrome P450 | 8 of 68 | 1.52 | 2.13e-08 |
| mmu00983 | Drug metabolism - other enzymes | 8 of 89 | 1.4 | 1.23e-07 |
| mmu01524 | Platinum drug resistance | 7 of 77 | 1.41 | 1.04e-06 |
| mmu05418 | Fluid shear stress and atherosclerosis | 8 of 141 | 1.21 | 2.61e-06 |
| mmu05169 | Epstein-Barr virus infection | 9 of 214 | 1.08 | 3.78e-06 |
| mmu05160 | Hepatitis C | 8 of 158 | 1.16 | 4.69e-06 |
| mmu05164 | Influenza A | 8 of 167 | 1.13 | 6.34e-06 |
| mmu05225 | Hepatocellular carcinoma | 8 of 168 | 1.13 | 6.34e-06 |

Biological Process (Gene Ontology)

| GO-term | description | count in network | strength | false discovery rate |
|---|---|---|---|---|
| GO:0051707 | Response to other organism | 38 of 1145 | 0.97 | 1.13e-23 |
| GO:0044419 | Interspecies interaction between organisms | 39 of 1309 | 0.93 | 2.07e-23 |
| GO:0051607 | Defense response to virus | 21 of 170 | 1.54 | 1.36e-22 |
| GO:0098542 | Defense response to other organism | 28 of 775 | 1.01 | 1.20e-17 |
| GO:0009605 | Response to external stimulus | 41 of 2199 | 0.72 | 1.20e-17 |
| GO:0002252 | Immune effector process | 22 of 435 | 1.16 | 3.61e-16 |
| GO:0045087 | Innate immune response | 23 of 558 | 1.07 | 3.30e-15 |
| GO:0006955 | Immune response | 28 of 979 | 0.91 | 3.30e-15 |
| GO:0006952 | Defense response | 29 of 1133 | 0.86 | 1.04e-14 |
| GO:0002376 | Immune system process | 33 of 1842 | 0.7 | 7.96e-13 |
| GO:0009617 | Response to bacterium | 21 of 660 | 0.95 | 1.28e-11 |
| GO:0006950 | Response to stress | 38 of 3045 | 0.55 | 1.76e-10 |
| GO:0050896 | Response to stimulus | 55 of 6908 | 0.35 | 1.26e-09 |
| GO:0035456 | Response to interferon-beta | 7 of 43 | 1.66 | 3.45e-07 |
| GO:0006749 | Glutathione metabolic process | 7 of 53 | 1.57 | 1.22e-06 |

**Sup Fig 3-1: Age-specific effects of C-Myc expression in old mouse hepatocytes.**
A) AAV-TBG-mApple expression in isolated hepatocytes ex-vivo after 2-weeks infection. B) Heatmap of methylation changes induced by Myc or mApple in young and old liver (N=4/cohort). C) Principal component analysis of RNAseq of young and old mice given AAV-TBG- mApple, c-Myc or Ccnd1. D) Number of DEGs for oncogenes by age along with overlapping genes. E) Heatmap of liver identity gene expression by age and treatment. F) Heatmap of old Myc only DEG. G) KEGG and gene ontology of old only Myc DEGs.

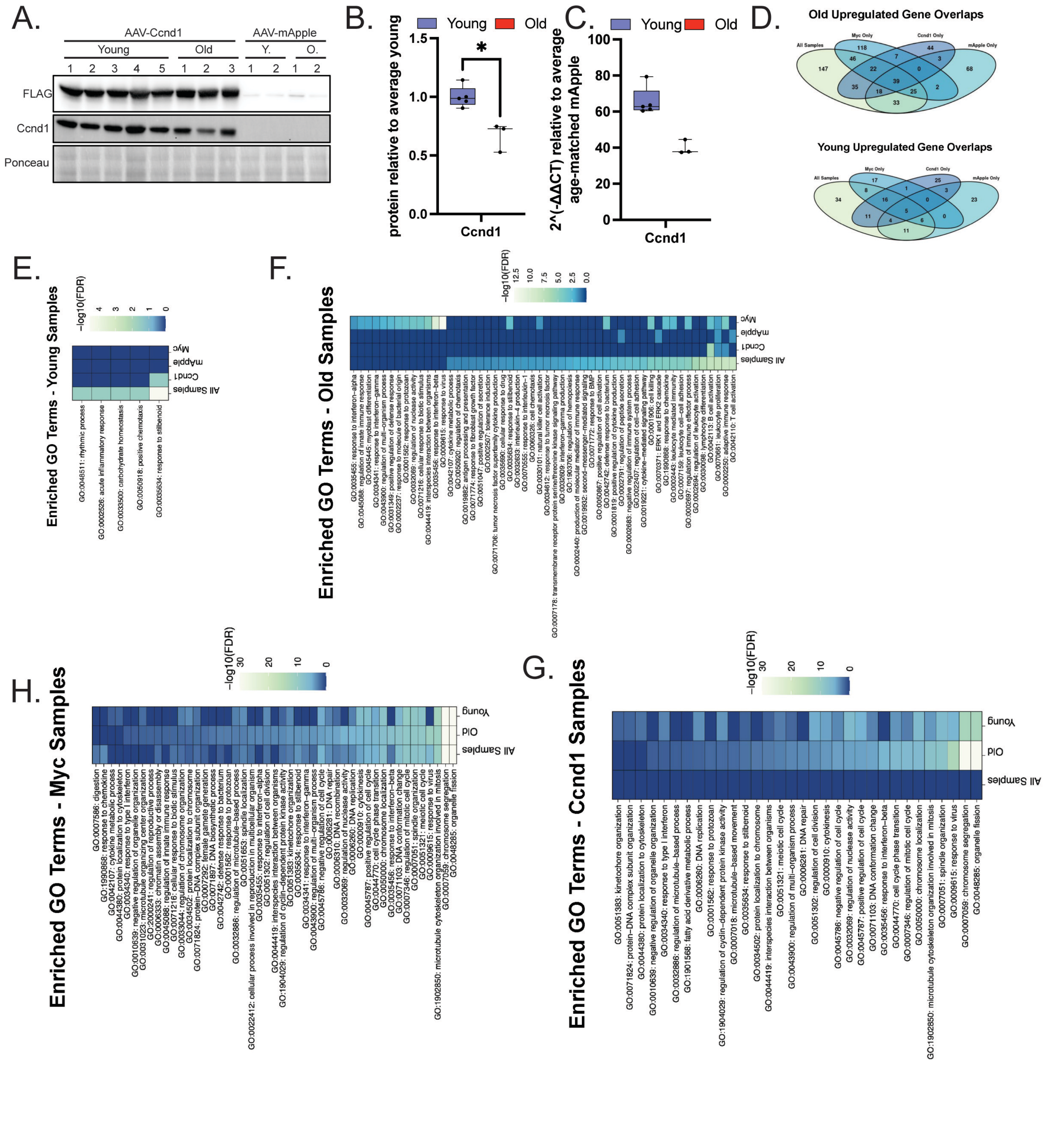

**Sup Fig 3-2: transcriptomic effects of AAV-TBG-Ccnd1 expression in young and old mouse hepatocytes.**
A) Western blot of Ccnd1 expression in young and old infected for 30-days AAV-TBG-Ccnd1/3xFLAG. B) Quantitation of western blot for Ccnd1 expression in young and old. C) qPCR analysis of Ccnd1 expression in young and old AAV-TBG-Ccnd1 treated mice. D) shows overlap of DEGs by age for mApple, Myc and Ccnd1 expression. E-G) Enriched gene ontology pathways for young and old mice comparing indicated treatments. Statistical analysis represented using one-way ANOVA with post hoc Tukey test in which asterisk (*) denotes $p < 0.05$.

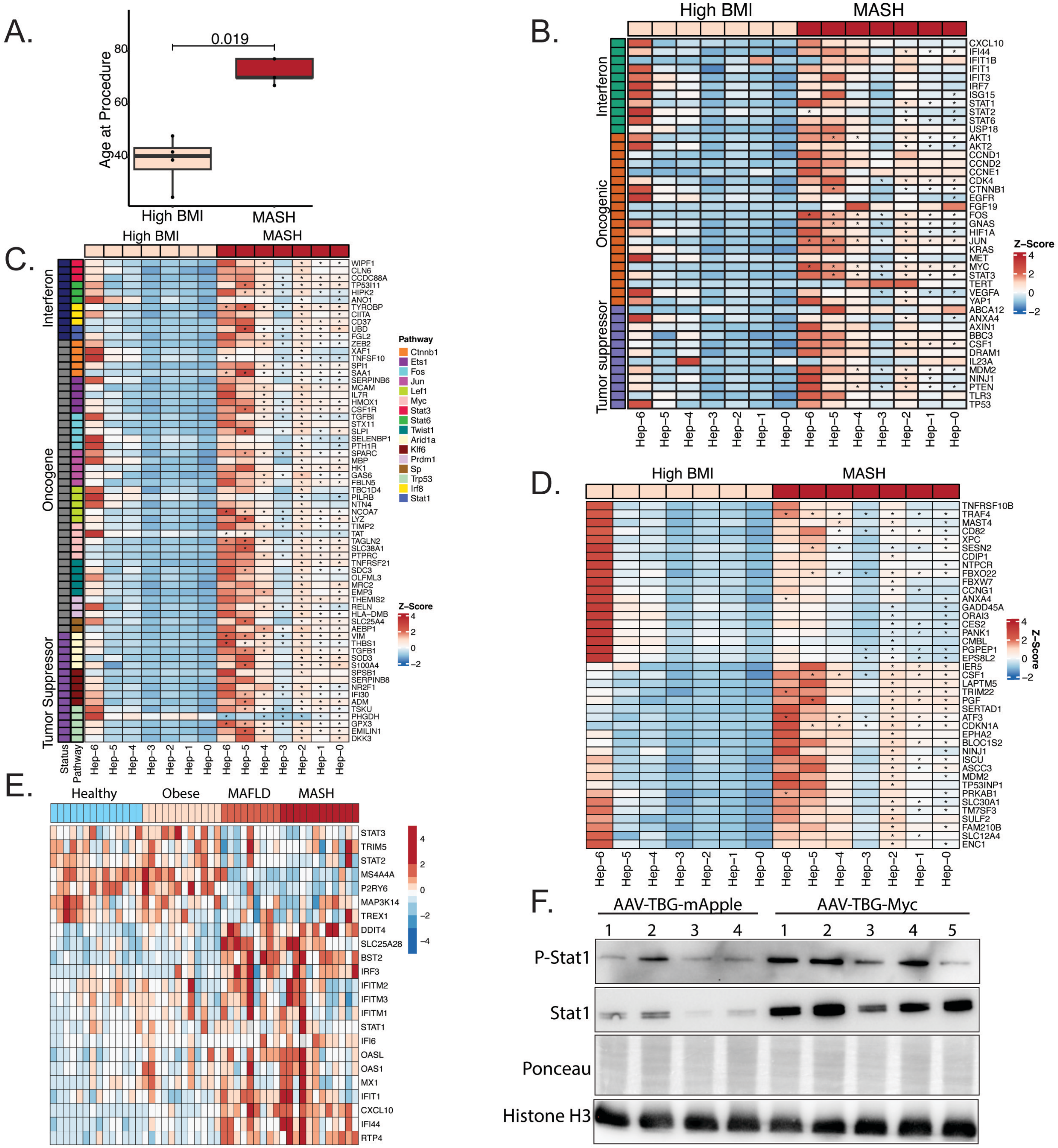


**Sup Fig 3-3: Elevation of IFN, oncogenic and tumor suppressor signaling with liver disease progression and age.** A) Age distribution by condition with statistics from Visium human data. B) Heatmap of Visium data representing expression of ISGs, oncogenes and tumor suppressors. C) Expression of IPA predicted downstream targets of interferon signaling, tumor suppressors and oncogenes. D) Expression of p53 target genes in different hepatocyte clusters of high BMI and MASH samples. E) Heatmap of human healthy, obese, MAFLD and MASH from published data set. F) Western blot of P-S727 Stat1, total Stat1 in AAV-TBG-mApple vs AAV-TBG-Myc treated old mice. Statistical analysis represented using one-way ANOVA with post hoc Tukey test in which asterisk (*) denotes $p < 0.05$.

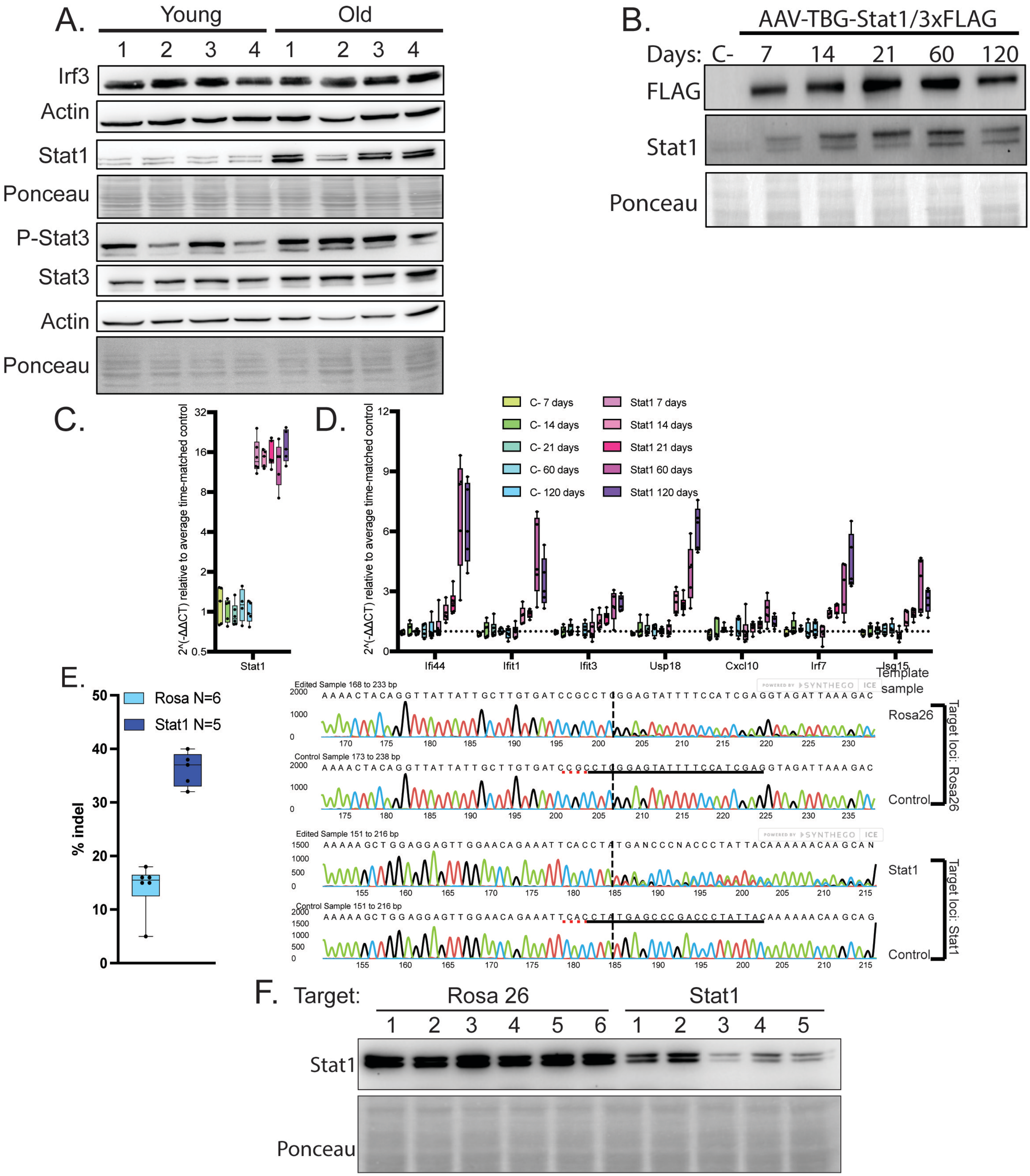


**Sup Fig 4-1: Elevated Stat1 is sufficient and necessary for ISG upregulation.**
A) Western blot of Irf3, Stat3, Phos-Stat3, Stat1 in isolated hepatocytes of young (5-MO, N=4) and old (22-MO, N=4) mice. B) Representative western blot for Stat1 and FLAG expression in 120-day time course of AAV-TBG-Stat/3xFLAG vs AAV-TBG-MCS. C) qPCR expression of Stat1 for full time-course with AAV-TBG-Stat1/3xFLAG vs AAV-TBG-MCS D) ISG expression for full time-course with AAV-TBG-Stat1/3xFLAG vs AAV-TBG-MCS (N=5/cohort). Note all ISGs are significantly elevated beginning day 14 and continue through the remainder of the time-course when treated with AAV-TBG-Stat1/3xFLAG. E) SaCas9 sgRosa (N=6) vs sgStat1 (N=5) ko % indels and representative sanger sequence after 21-days treatment with AAV in old (25-MO) mice. F) Western blot of Stat1 in AAV-TBG-saCas9 with guides targeting Rosa26 (N=6) or Stat1 (N=5) for 21-days in old (25-MO) mice.

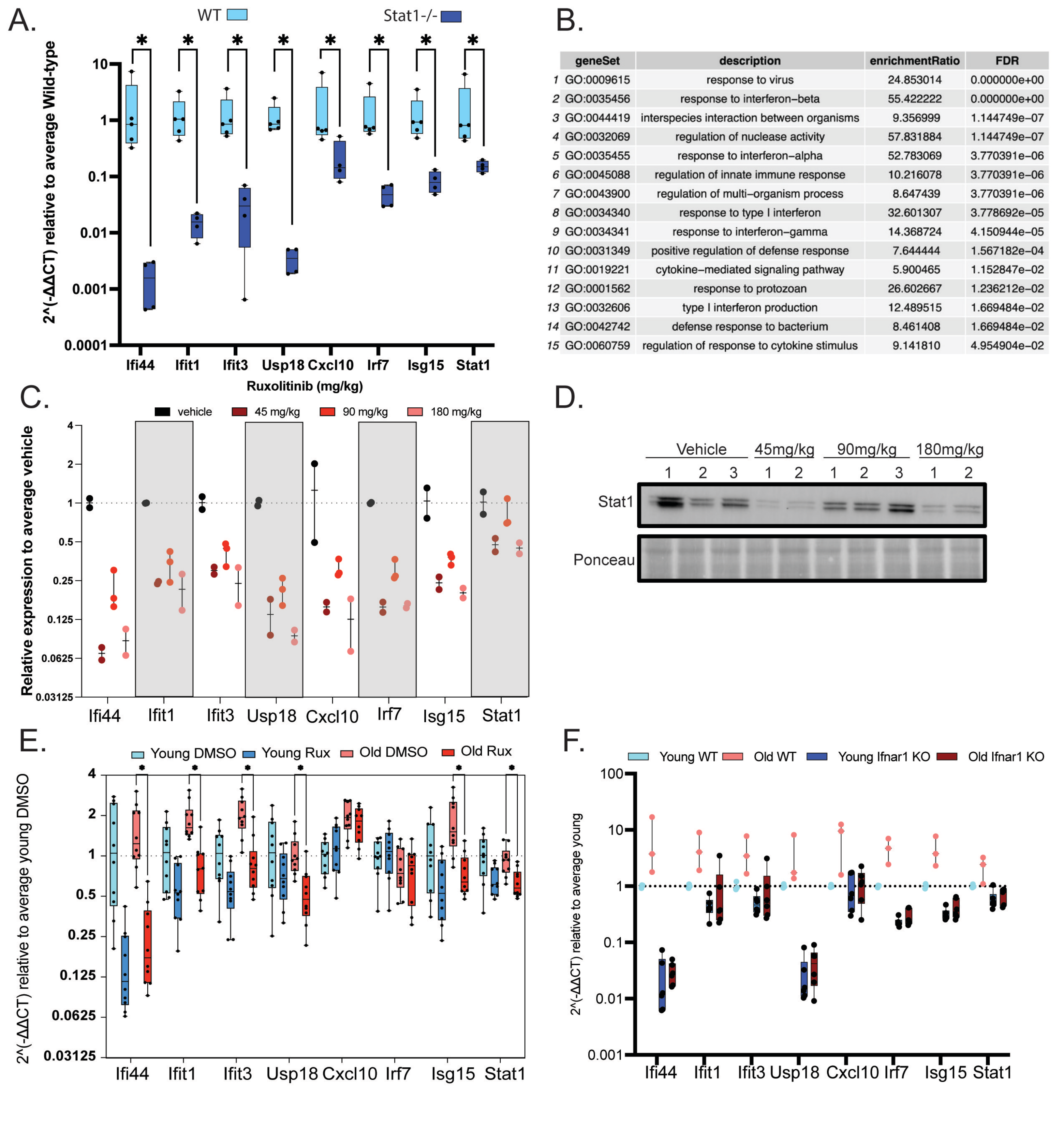


| | geneSet | description | enrichmentRatio | FDR |
|---|---|---|---|---|
| 1 | GO:0009615 | response to virus | 24.853014 | 0.000000e+00 |
| 2 | GO:0035456 | response to interferon−beta | 55.422222 | 0.000000e+00 |
| 3 | GO:0044419 | interspecies interaction between organisms | 9.356999 | 1.144749e−07 |
| 4 | GO:0032069 | regulation of nuclease activity | 57.831884 | 1.144749e−07 |
| 5 | GO:0035455 | response to interferon−alpha | 52.783069 | 3.770391e−06 |
| 6 | GO:0045088 | regulation of innate immune response | 10.216078 | 3.770391e−06 |
| 7 | GO:0043900 | regulation of multi−organism process | 8.647439 | 3.770391e−06 |
| 8 | GO:0034340 | response to type I interferon | 32.601307 | 3.778692e−05 |
| 9 | GO:0034341 | response to interferon−gamma | 14.368724 | 4.150944e−05 |
| 10 | GO:0031349 | positive regulation of defense response | 7.644444 | 1.567182e−04 |
| 11 | GO:0019221 | cytokine−mediated signaling pathway | 5.900465 | 1.152847e−02 |
| 12 | GO:0001562 | response to protozoan | 26.602667 | 1.236212e−02 |
| 13 | GO:0032606 | type I interferon production | 12.489515 | 1.669484e−02 |
| 14 | GO:0042742 | defense response to bacterium | 8.461408 | 1.669484e−02 |
| 15 | GO:0060759 | regulation of response to cytokine stimulus | 9.141810 | 4.954904e−02 |

**Sup Fig 4-2: Jak/Stat1 inhibtion ameliorates ISG expression.**
A) qPCR of ISGs of Stat1-/- (5-MO, N=4) mice vs age-matched wildtype (N=5) mice. B) Gene ontology overrepresentation analysis of RNAseq of old (24-MO) mice that received either combination of anti-IrnaR1 and anti-Ifn⍰R1 blockade or isotype control blockade for 21-days C) qPCR of ISGs after 5-day treatment with ruxolitinib at 45, 90 and 180mg/kg given oral gavage 2x daily in old (24-MO N=2 or 3/cohort) mice. D) Western blot of Stat1 for dose response associated with ruxolitinib. E) qPCR of hepatocytes isolated from young (6-MO) and old (23-MO) mice cultured ex Vivo treated with 2uM ruxolitinib or DMSO for 24-hours (N=10/cohort). F) qPCR of ISGs for Ifnar1-/- (N=6/age) young (7-MO) and old (21-MO) mice age matched vs wildtype mice (N=3/age). Statistical analysis represented using Mann-Whitney in which asterisk (*) denotes $p < 0.05$.

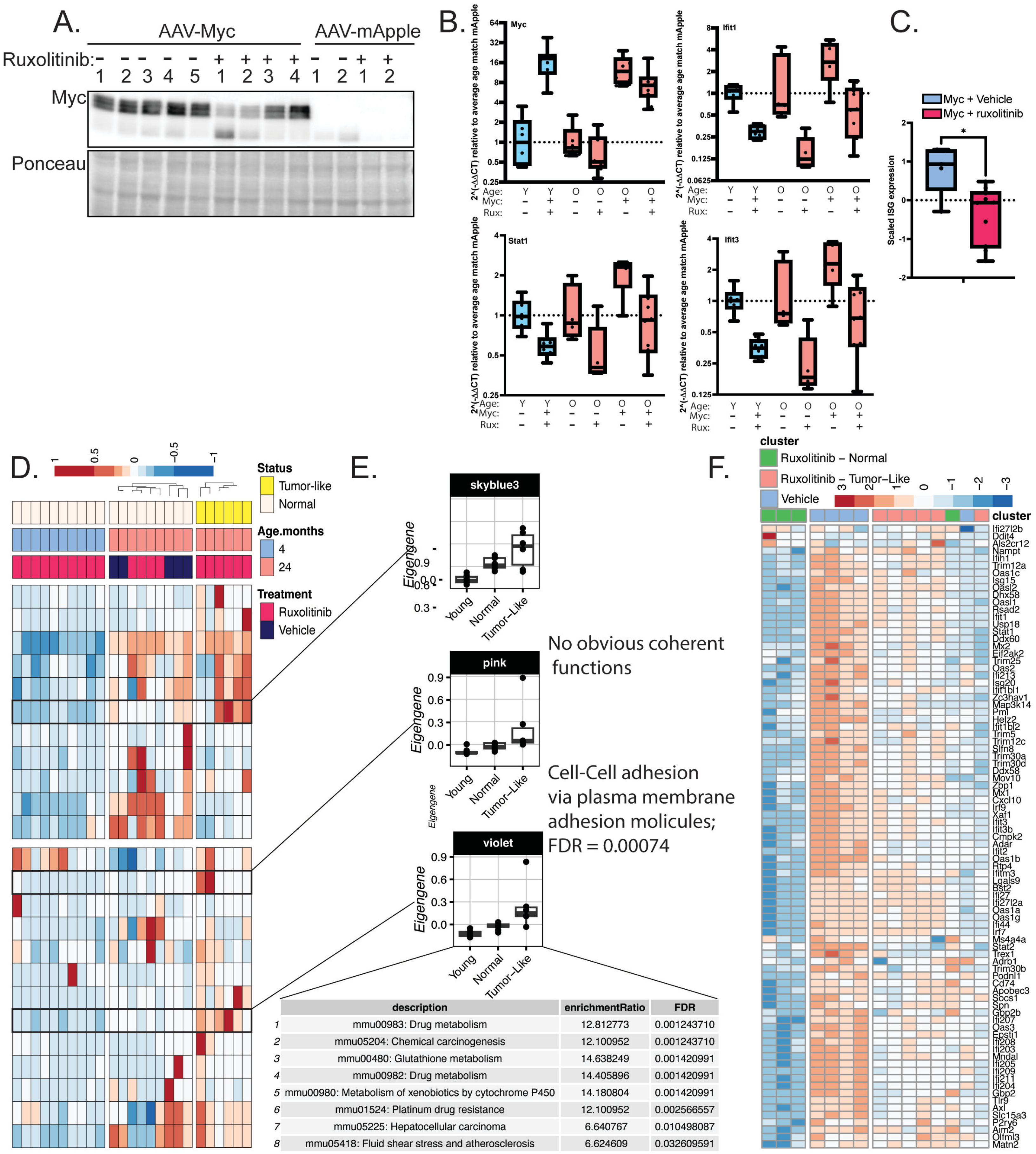


| | description | enrichmentRatio | FDR |
|---|---|---|---|
| 1 | mmu00983: Drug metabolism | 12.812773 | 0.001243710 |
| 2 | mmu05204: Chemical carcinogenesis | 12.100952 | 0.001243710 |
| 3 | mmu00480: Glutathione metabolism | 14.638249 | 0.001420991 |
| 4 | mmu00982: Drug metabolism | 14.405896 | 0.001420991 |
| 5 | mmu00980: Metabolism of xenobiotics by cytochrome P450 | 14.180804 | 0.001420991 |
| 6 | mmu01524: Platinum drug resistance | 12.100952 | 0.002566557 |
| 7 | mmu05225: Hepatocellular carcinoma | 6.640767 | 0.010498087 |
| 8 | mmu05418: Fluid shear stress and atherosclerosis | 6.624609 | 0.032609591 |

**Sup Fig 5-1: Heterogenous transcriptional effects of Myc + ruxolitinib in old mice have patterns of transformation.** A) Western blot of Myc expression in young and old mouse liver for indicated treatment. B) qPCR Myc and indicated ISGs. C) Boxplot of normalized Z-score for ISG expression comparing old + Myc vehicle and old + Myc + ruxolitinib samples. D) WGCNA analysis of all Myc treated mouse samples used to determine different clusters represented in Myc + ruxolitinib treated samples. E) KEGG pathway analysis of WGCNA modules associated with tumor-like samples. F) Heatmap of ISG expression in old AAV-TBG-Myc ruxolitinib or vehicle treated mice. Statistical analysis represented using one-way ANOVA with post hoc Tukey test in which asterisk (*) denotes $p < 0.05$, (**) $p < 0.01$, (***) $p < 0.001$ and (****) $p < 0.0001$.

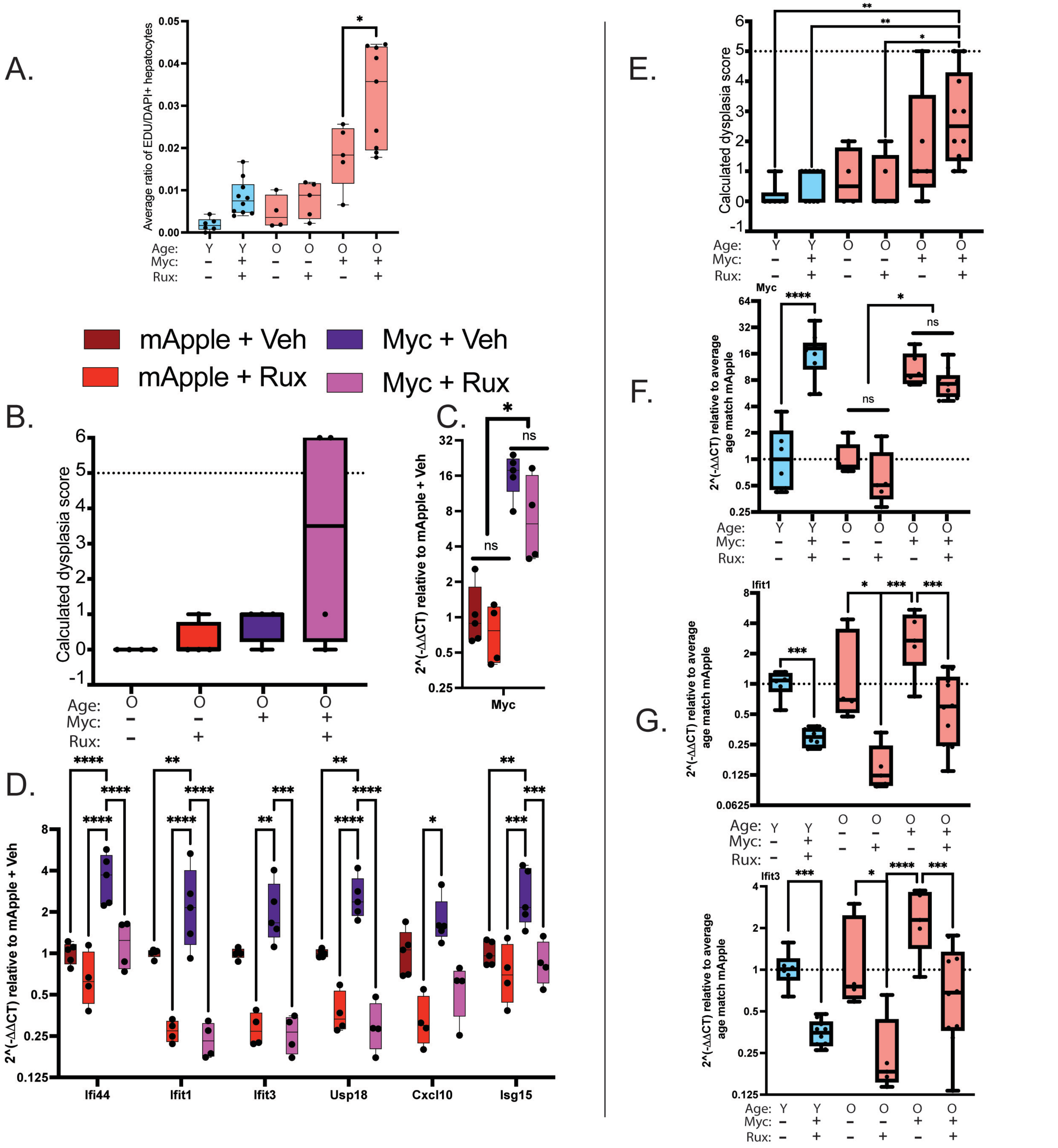


**Sup Fig 5-2: Two independent experiments show elevated dysplasia or tumorigenesis only in old mice with Myc and ruxolitinib.**
Two independent experiments examining response to AAV-TBG-mApple or Myc in: A) quantitation of average ratio of EdU+/DAPI hepatocytes as observed by immunofluorescence. B) Calculated dysplasia score for experiment 1. C) qPCR of Myc expression. D) ISGs by treatment in experiment 1. E) Calculated dysplasia score for experiment 2. F) qPCR of Myc expression G) qPCR of representative ISGs by treatment. Statistical analysis represented using one-way ANOVA with post hoc Tukey test in which asterisk (*) denotes $p < 0.05$, (**) $p < 0.01$, (***) $p < 0.001$ and (****) $p < 0.0001$.

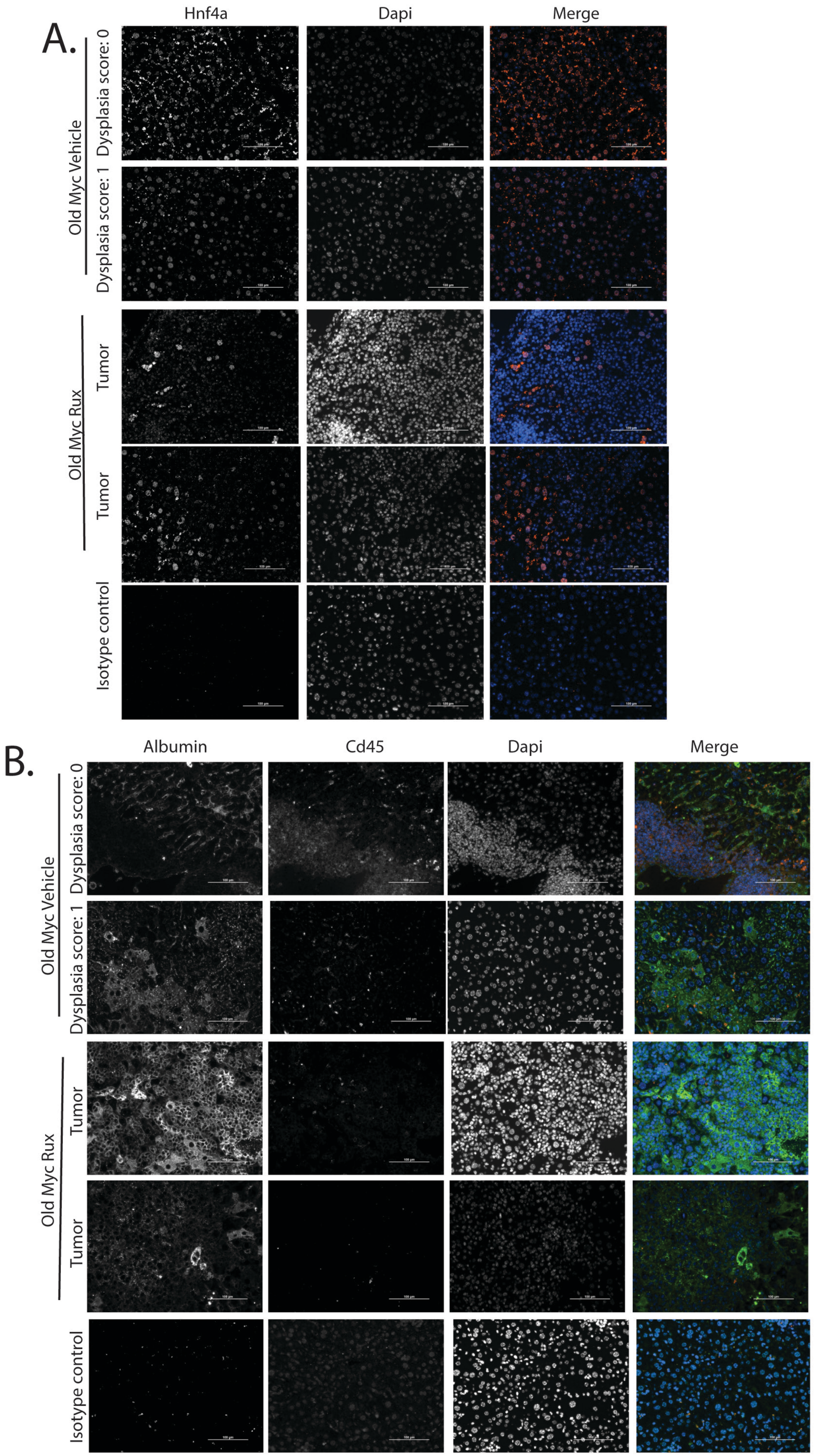


**Sup Fig 5-3: Immunofluorescence images of transformed regions within old mice given Myc and ruxolitinib examining markers of hepatocytes.**
A) Immunofluorescence old AAV-TBG-Myc vehicle vs AAV-TBG-Myc ruxolitinib transformed samples along with indicated dysplasia score for markers of hepatocytes: A) Hnf4a B) Albumin and immune cell marker Cd45.
